# *Pf*MORC protein regulates chromatin accessibility and transcriptional repression in the human malaria parasite, *Plasmodium falciparum*

**DOI:** 10.1101/2023.09.11.557253

**Authors:** Z Chahine, M Gupta, T Lenz, T Hollin, S Abel, CAS Banks, A Saraf, J Prudhomme, S Bhanvadia, L Florens, KG Le Roch

## Abstract

The environmental challenges the human malaria parasite, *Plasmodium falciparum*, faces during its progression into its various lifecycle stages warrant the use of effective and highly regulated access to chromatin for transcriptional regulation. Microrchidia (MORC) proteins have been implicated in DNA compaction and gene silencing across plant and animal kingdoms. Accumulating evidence has shed light into the role MORC protein plays as a transcriptional switch in apicomplexan parasites. In this study, using CRISPR/Cas9 genome editing tool along with complementary molecular and genomics approaches, we demonstrate that *Pf*MORC not only modulates chromatin structure and heterochromatin formation throughout the parasite erythrocytic cycle, but is also essential to the parasite survival. Chromatin immunoprecipitation followed by deep sequencing (ChIP-seq) experiments suggest that *Pf*MORC binds to not only sub-telomeric regions and genes involved in antigenic variation but may also play a role in modulating stage transition. Protein knockdown experiments followed by chromatin conformation capture (Hi-C) studies indicate that downregulation of *Pf*MORC impairs key histone marks and induces the collapse of the parasite heterochromatin structure leading to its death. All together these findings confirm that *Pf*MORC plays a crucial role in chromatin structure and gene regulation, validating this factor as a strong candidate for novel antimalarial strategies.

## Introduction

Malaria is a mosquito-borne infectious disease that is caused by protozoan parasites of the genus *Plasmodium*. Among the five human-infecting species, *Plasmodium falciparum* is the deadliest, with over 619,000 deaths in 2023(*1*). To adapt to extreme environmental challenges, *P. falciparum* possesses unique strategies that direct the tightly coordinated changes in gene expression and control transcriptional switching in genes encoded by multigene families to ensure antigenic variation and immune evasion. Changes in gene expression throughout the parasite life cycle are controlled by a surprisingly low repertoire of transcription factors (TFs) encoded in the *Plasmodium* genome (*2–9*). The 27 apicomplexan APETALA2 (ApiAP2) DNA-binding proteins are the only well documented parasite TFs known to contribute to the modulation of gene expression throughout various stages of the parasite’s development(*3, 5–8, 10–13*). Since the discovery of the AP2 gene family(*3*), evidence has alluded to their role as master regulators of gene expression throughout transitory phases of the parasite life cycle. The most well documented are the gametocyte specific TFs (AP2-G, AP2-G2)(*7, 14*) with subsequent discovery of those associated with transition to sexual differentiation(*13*) as well as sporozoite (AP2-SP) and liver stages (AP2-L) (*6, 10*). The *Plasmodium* genome, however, encompasses well over 5,000 protein-coding genes suggesting there are most likely other molecular components responsible for gene expression. It is believed that, to offset the relatively low TF range, *Plasmodium* has evolved additional mechanisms regulating gene expression including mechanisms that use epigenetics factors, RNA binding proteins or regulate chromatin structure (*15–20*). All together these components work in combination to regulate the dynamic organization of DNA and the cascade of gene expression required for the parasite life cycle progression(*21–28*). Despite their importance, the identification and functional characterization of regulatory players controlling chromatin remains a challenge in this intractable organism. However, with the advent of sensitive technologies capable of capturing important chromatin associated regulatory complexes such as chromatin immunoprecipitation (ChIP), chromatin conformation capture Hi-C technologies, and chromatin enrichment for proteomics (ChEP), we can now solve important facets of molecular components controlling epigenetics and chromatin structure.

Microrchidia (MORC) belong to a highly conserved nuclear protein superfamily with widespread domain architectures that link MORCs with signaling-dependent chromatin remodeling and epigenetic regulation across plant and animal kingdoms, including the apicomplexan parasites(*29–32*). In all organisms, MORC contains several domains that form a catalytically active ATPase. Apicomplexan MORC ATPases are encircled by Kelch-repeat β-propellers as well as a CW-type zinc finger domain functioning as a histone reader(*33*). In higher eukaryotes, MORCs were first identified as epigenetic regulators and chromatin remodelers in germ cell development. Currently, these proteins are shown to be involved in various human diseases including cancers and are expected to serve as important biomarkers for diagnosis and treatment(*34*). In the apicomplexan parasite, *Toxoplasma gondii*, *Tg*MORC was shown to recruit the histone deacetylase HDAC3 to particular genome loci to regulate chromatin accessibility, restricting sexual commitment (*33, 35*). *Tg*MORC-depleted cells also resulted in change in gene expression with up regulation of secondary AP2 factors and a significant shift from asexual to sexual differentiation(*32, 33, 36–38*). Having multiple homologs with *T. gondii*, *Plasmodium* AP2 protein conservation is primarily restricted to their AP2 DNA-binding domains (*39, 40*). Recently, ChEP experiments done in *P. falciparum* identified *Pf*MORC as one of the highest enriched chromatin-bound proteins at different stages of the parasite intraerythrocytic development cycle (IDC)(*28*). *Pf*MORC was also detected at a relatively high level throughout the parasite life cycle, including sporozoites and liver stages(*33, 41*), and has been identified in several protein pull down experiments as targeting both AP2 TFs and epigenetic factors(*32, 33, 38, 42–44*). Immunoprecipitation experiments demonstrated that *Pf*MORC seems to interact with AP2-G2(*32*), a TF that plays a critical role in the maturation of gametocytes. However, the genome wide distribution and exact function of this protein throughout the parasite life cycle remained elusive.

In this study, we apply CRISPR/Cas9 genomic editing technologies to determine the function, genome distribution, and indispensability of *Pf*MORC throughout the parasite’s IDC. Immunoprecipitation of an HA-tagged parasite line validates the role of *Pf*MORC in heterochromatin structure maintenance. Using downregulation of *Pf*MORC induced through TetR-DOZI system, we demonstrate the functional significance of *Pf*MORC in heterochromatin stability and gene repression. Immunofluorescence based assays and ChIP-seq experiments show that *Pf*MORC localizes to heterochromatin clusters at or near *var* genes with significant overlap with well-known signatures of the parasite heterochromatin post-translational modification (PTM) H3K9-trimethylation (H3K9me3) marks. When *Pf*MORC was downregulated, level of H3K9me3 was detected at a lower level, demonstrating a possible role of this protein in epigenetic regulation and gene repression. Finally, Hi-C analyses demonstrate that downregulation of *Pf*MORC results in significant dysregulation of chromatin architectural stability resulting in parasite death. All together our work provides significant insight into the role of *Pf*MORC in the maintenance of the pathogens’ epigenetics and chromosomal architectural integrity and validates this protein as a promising target for novel therapeutic interventions.

## Results

### Generation of *P*fMORC-HA transgenic line shows nuclear localization in heterochromatin clusters

To characterize the role of *Pf*MORC in *P. falciparum*, we applied CRISPR/Cas9 genome editing tool to add a 3X-HA tag at the C-terminal coding region of the *Pfmorc* locus (PF3D7_1468100) in an NF54 line(*45*) (**Fig. 1a, Sup. data 1**). Recovered transgenic parasites were cloned using limiting dilution and the correct incorporation of the tag into the parasite genome was validated via PCR and whole genome sequencing (WGS) (**Fig. 1b, Sup. data 2**). WGS analysis confirmed the presence of the HA tag at the expected locus with no obvious off-target effect but uncovered an additional nonsense mutation in the *gametocyte development protein 1* (*gdv1*) gene (PF3D7_0935400) at amino acid position 561 out of 599 total. Mutations in *gdv1* have been detected in the past and seem to emerge relatively often, indicating a fitness benefit in the in *vitro* culture system used in the laboratory(*8*). The involvement of GDV1 as essential in sexual commitment is well substantiated and further study involving the role of *Pf*MORC in sexual commitment could not be fully addressed(*7, 8, 14, 46–48*)(**Sup. data 2**). Despite this finding, we were able to design experiments to determine the role of *Pf*MORC throughout the IDC as no other major detrimental mutations were detected in the genome. We first validated *Pf*MORC protein expression through Western blot analysis (**Fig. S1**). Regardless of potential protein degradation, results showed an expected band size within the ∼290 kDa range that was absent in NF54 control compared to our tagged line (**Fig. S1**). Immunofluorescence (IFA) analysis of intracellular parasites revealed that *Pf*MORC is localized in the nucleus throughout the parasite IDC in punctate patterns (**Fig. 1c**). A single foci per parasite was detected in the nuclear periphery at the ring stage. An increased number of foci with a more diffuse signal could be detected as the number of DNA copies increased at the schizont stage. Moreover, punctuates were shown to have strong colocalization signals with histone H3K9me3 marks present at the different parasite developmental stages analyzed. This was most evident during the early ring stage of the parasite IDC where the chromatin organization is more compact with a single DNA copy. Altogether, our findings indicated that *Pf*MORC is a true nuclear protein that is most likely associated with *P. falciparum* heterochromatin cluster(s).

**Fig. 1.**
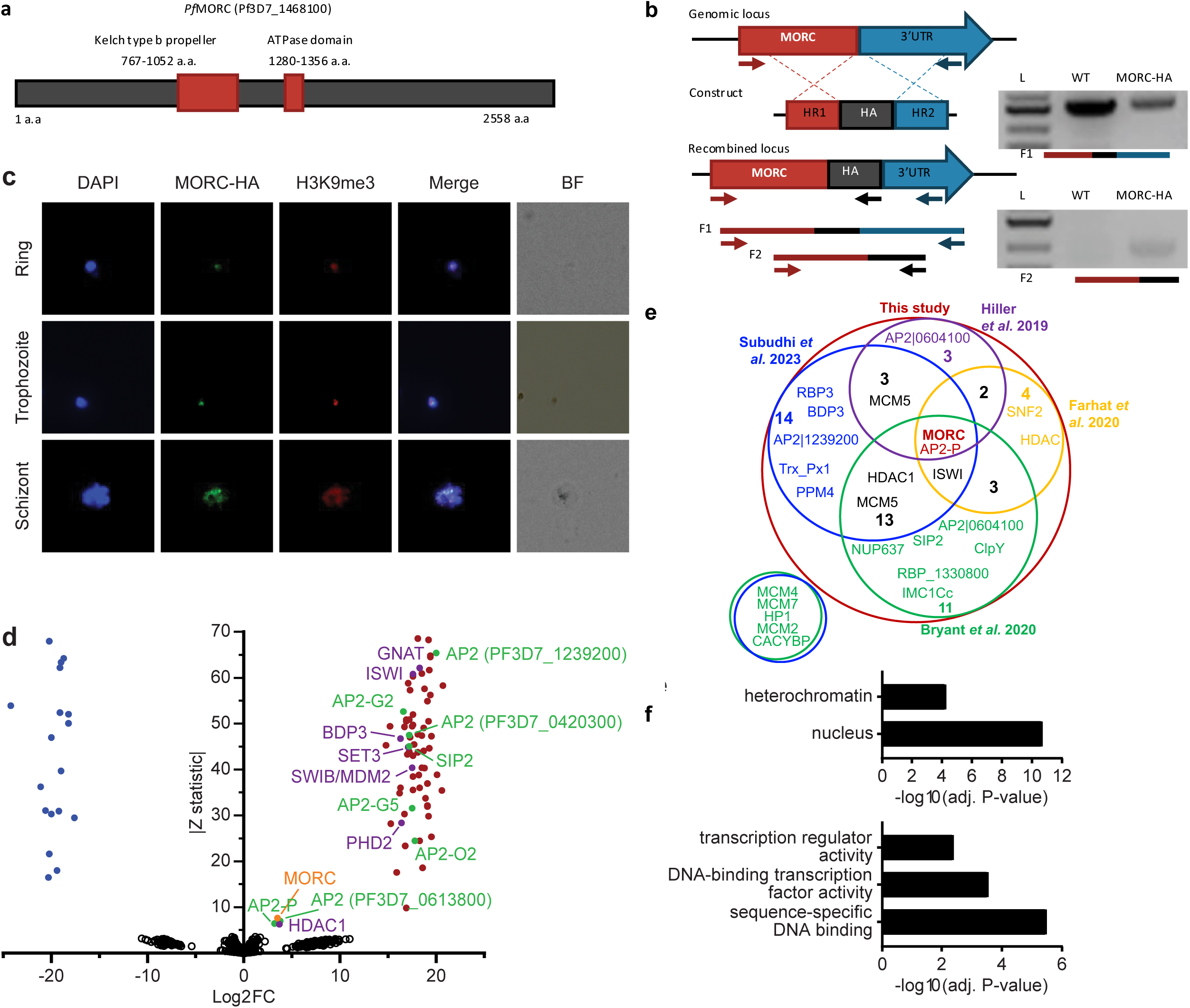
*Pf*MORC-HA is associated with heterochromatin. **(a)** Illustration of the *Pf*MORC containing domains including Kelch type b propeller and ATPase domains using InterProScan. **(b)** Design strategy applied for *Pf*MORC C-terminal HA tagging. PCR amplification of the genomic C-terminus end of *Pfmorc* region extending towards the 3’UTR (F1) as well as extension from the C-terminus towards the HA flanking sequence (F2) verifies the correct insertion site. NF54 genomic DNA was used as negative control. **(c)** IFA experiment: *Pf*MORC foci ^22^ expressing co-localization with H3K9me3 marks (red). Cell nuclei are stained with DAPI (blue). BF: brightfield **(d)** Protein immunoprecipitation: Significance plot representing *Pf*MORC interactome recovered through immunoprecipitation followed by mass spectrometry (IP-MS). Graph list MORC (orange) bindings partners of highest affinity associated with TF regulation ^22^ ^22^ and chromatin remodelers, erasers, and writers (purple). Proteins enriched in the *Pf*MORC-HA samples compared with controls were filtered with log_2_ FC ≥ 2 and Z statistic > 5. **(e)** Venn diagram representing overlapping proteins identified among five publications. Values represent the total number of significant proteins identified as overlapping between two subsets. **(f)** Gene ontology enrichment analysis of the significantly enriched proteins. The top 2 terms of Cellular Component (top) and top 3 terms of Molecular Function (bottom) are represented as -Log_10_ (adjusted P-value) (Fisher’s exact test with Bonferroni adjustment).

### *Pf*MORC interacting partners include proteins involved in heterochromatin maintenance, transcription regulation and chromatin remodeling

To confirm the association of *Pf*MORC with proteins involved in heterochromatin cluster(s), we performed immunoprecipitation (IP) followed by mass-spectrometry (MS) analysis using mature stages of *P*fMORC-HA and parental line as control. MORC was detected with highest peptides (97 and 113) and spectra (1041 and 1177) counts compared to control conditions (5 and 7 peptides; 16 and 43 spectra) confirming the efficiency of our pull-down. However, considering the relatively large size of the MORC protein (295kDa) and its weak detection in the control, the log_2_ FC and Z-statistic after normalization are minimal when compared to smaller proteins that were not identified in the control samples. To define a set of *Pf*MORC associated proteins, we used the QPROT statistical framework(*49*) to compare proteins detected in *Pf*MORC samples and controls. We identified 73 *Pf*MORC associated proteins (log_2_ FC ≥ 2 and Z statistic > 5) (**Fig. 1d, Sup. data 3**). Additional chromatin remodeling proteins were detected such as ISWI chromatin-remodeling complex ATPase (PF3D7_0624600) and SWIB/MDM2 domain-containing protein (PF3D7_0611400), as well as the epigenetic readers BDP3 (PF3D7_0110500) and PHD2 (PF3D7_1433400)(*50*). We also detected a significant enrichment of histone erasers and writers such as a N-acetyltransferase (PF3D7_1020700), HDAC1 (PF3D7_0925700), and SET3 (PF3D7_0827800). Several of which have been detected to interact with MORC in previous studies including the AP2-G5, HDAC1, ELM2 and ApiAp2 proteins(*33, 42–44*) (**Fig. 1e**). Notably, we observed that many of the detected proteins were enriched at levels comparable to those of the bait protein such as AP2-P and HDAC1. This may be a result of proteins forming complexes in a one-to-one ratio, leading to similar enrichment levels. Notably, two of these three proteins have been reported to interact with MORC in several studies, further supporting a strong interaction between them. Gene Ontology (GO) enrichment analysis revealed that these proteins are nuclear and are associated with heterochromatin, DNA binding, and transcription factor activity (**Fig. 1f**). Detailed analysis showed the detection of SIP2, involved in heterochromatin formation and chromosome end regulation(*51*), and 8 AP2 transcription factors, including AP2-G2, AP2-O2, AP2-G5. Among the other AP2s detected, 5 were previously identified by ChIP-seq as *Plasmodium falciparum* AP2 Heterochromatin-associated Factors (*Pf*AP2-HFs)(*52*). Our findings, corroborated with several published works, indicate that *Pf*MORC interacts with multiple ApiAP2 TFs, chromatin remodelers, and epigenetic players associated with heterochromatin regions.

### Genome-wide distribution of *Pf*MORC displays functional roles in stage-specific gene silencing and antigenic variation

Given the multifunctional role of MORC proteins throughout apicomplexan parasites, we further explored the association of *Pf*MORC with chromatin accessibility and stage-specific gene regulation by assessing the genome-wide distribution of *Pf*MORC using chromatin immunoprecipitation followed by deep sequencing (ChIP–seq) in duplicate at the ring, trophozoite and schizont stages. Inspection of *Pf*MORC binding sites across each chromosome revealed a strong signal for telomeric and subtelomeric regions of the parasite genome, as well as the internal clusters of genes responsible for antigenic variation such as *var* and *rifin* (**Fig. 2a-c, Sup. data 4**). We found that while only 47% of reads mapped to antigenic genes in the ring stage, 95% of reads mapped to the same genes during the trophozoite stage and 82% during the schizont stage. Although a vast majority of the genome is actively transcribed during the trophozoite stage, these antigenic gene families are under tight regulatory control to ensure mutually exclusive expression and participate in immune evasion(*53, 54*). This could explain the disparity in the level of *Pf*MORC binding of the coding regions between the ring and trophozoite stages (**Fig. 2c, Fig. S2b**).

**Fig. 2.**
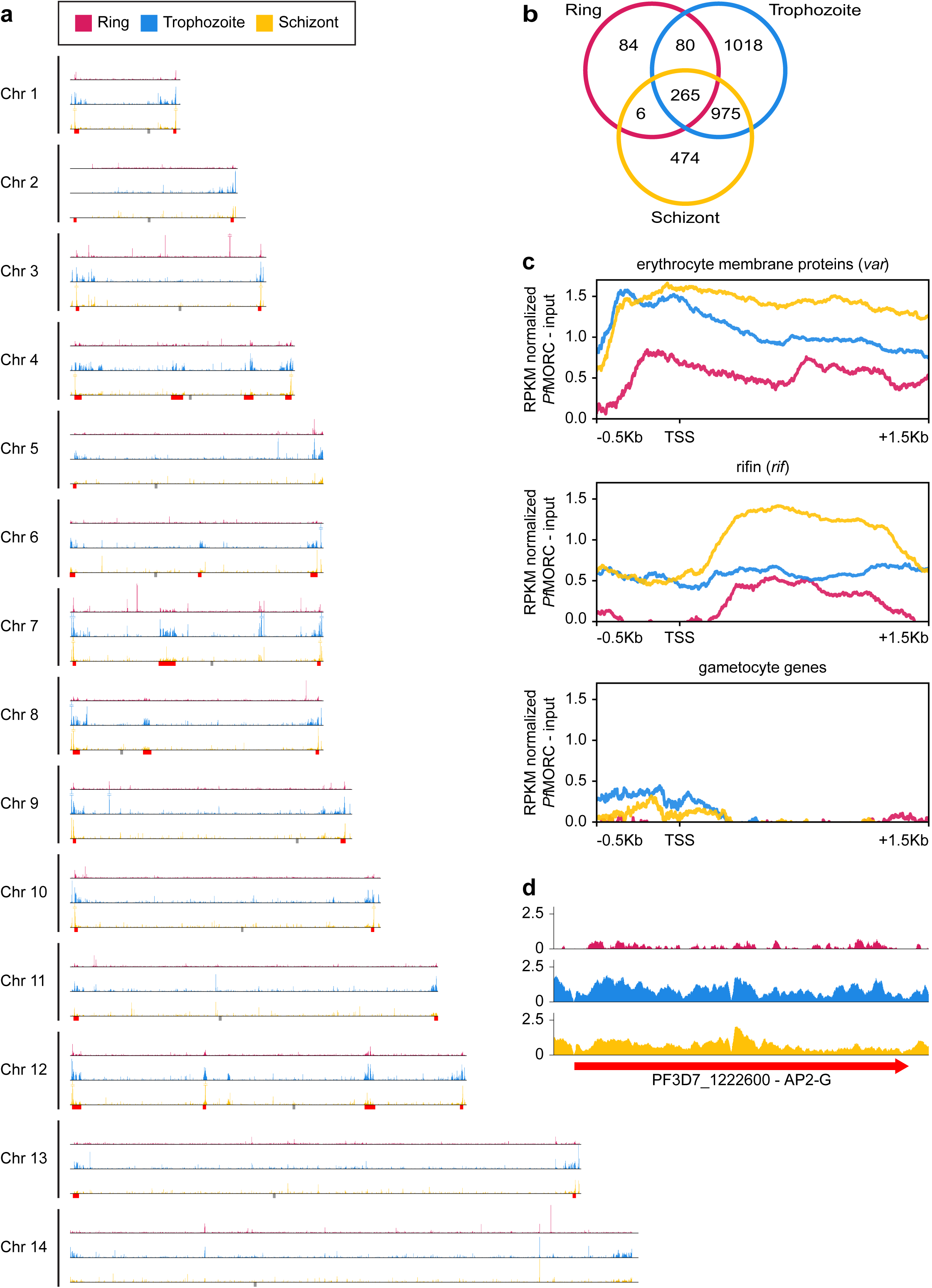
Genome wide distribution of *Pf*MORC proteins. **(a)** Chromosome distribution plots of *Pf*MORC binding showing a predisposition for subtelomeric and internal *var* gene regions (red). Each track is input subtracted, and per-million read count normalized before normalizing the track height to allow for direct comparison between stages. Grey boxes indicate the position of centromeres. **(b)** Overlap of called peaks between time points. **(c)** Profile plots showing *Pf*MORC coverage from 0.5 kb 5’ of the transcription start site (TSS) to 1.5 kb 3’ of the TSS in the ring, trophozoite, and schizont stages. Each plot includes per-million read count normalized coverage at 1 bp resolution for all genes within the *var* and *rifin* gene families as well as gametocyte-specific genes. **(d)** *Pf*MORC coverage of the gametocyte-specific transcription factor *ap2-g*.

We also evaluated *Pf*MORC binding of stage-specific gene families including gametocyte-related genes, and merozoite surface proteins (*msp*)(*55*). In trophozoite and schizont stages, gametocyte-associated genes contain a mean of <0.5 RPKM normalized reads per nucleotide of *Pf*MORC binding within their promoter region, whereas antigenic gene families such as *var* and *rifin* contain ∼1.5 and 0.5 normalized reads, respectively **(Fig. 2b)**. The difference is even greater within the gene body with gametocyte genes displaying almost no reads mapped more than 200 bp downstream of the TSS and antigenic gene families containing 0.5-1.5 RPKM normalized reads. However, the gametocyte-specific transcription factor AP2-G, known to be repressed during the IDC and required for sexual commitment, deviates from this trend and contains similar levels of *Pf*MORC binding to *var* genes **(Fig. 2c and S2c, d)**. These results indicate a major role of *Pf*MORC in controlling AP2-G and sexual differentiation. For some of the *msp* genes that are usually expressed later during the IDC to prepare the parasite for egress and invasion of new erythrocytes, *Pf*MORC binding was detected in the gene bodies at the trophozoite stage. We also detected a small switch in *Pf*MORC binding sites, moving from their gene bodies to their intergenic regions at the schizont stage (**Fig. S2d**). *Pf*MORC most likely moved away from the gene body to the regulatory regions surrounding the transcription start site (TSS) to guide RNA Polymerase and transcription factors and aid in activating expression of genes at the schizont stage that are crucial for egress and invasion. These results provide strong evidence for the direct effects that *Pf*MORC binding has on tightly controlled antigenic and stage-specific genes including crucial invasion and gametocyte genes.

### *P*fMORC is essential for *P. falciparum* survival

We next sought to confirm the functional relevance of *Pf*MORC protein in parasite survival. While partial *Pf*MORC knockdown using the glmS-ribozyme system had previously shown to have no significant effect on the parasite survival(*41*), another study using transposon mutagenesis (*piggyBac)* identified *Pf*MORC as likely essential(*56*). To resolve these conflicting results, we applied a complementary approach using the CRISPR/Cas9 gene editing strategy to incorporate an inducible TetR-DOZI system to knockdown (KD) *Pf*MORC(*57*) through administration of anhydrotetracycline (aTC) (**Fig. 3a**)(*45, 58*). The protein was also modified to include a C-terminal 3x-HA tag. Parental and transgenic clones were validated via PCR (**Fig. 3b**) and WGS to confirm the correct insertion of the inducible system (**Sup. data 2**) as well as the absence of major mutation that could explain some of the phenotypes observed. While results from our WGS validated our editing strategy without the detection of any obvious deleterious off-target effect, we identified a nonsense mutation in AP2-G gene (PF3D7_1222600) explaining our inability to obtain mature gametocytes. Protein expression and successful KD was confirmed via western blot with a decrease of ∼ 58% and ∼ 88.2% in *Pf*MORC expression at 24 hpi and 36 hpi respectively compared to their aTC supplemented conditions (**Fig. S3**). Downregulation of *Pf*MORC was detected significantly above the level observed by Singh and colleagues(*41*).

**Fig. 3.**
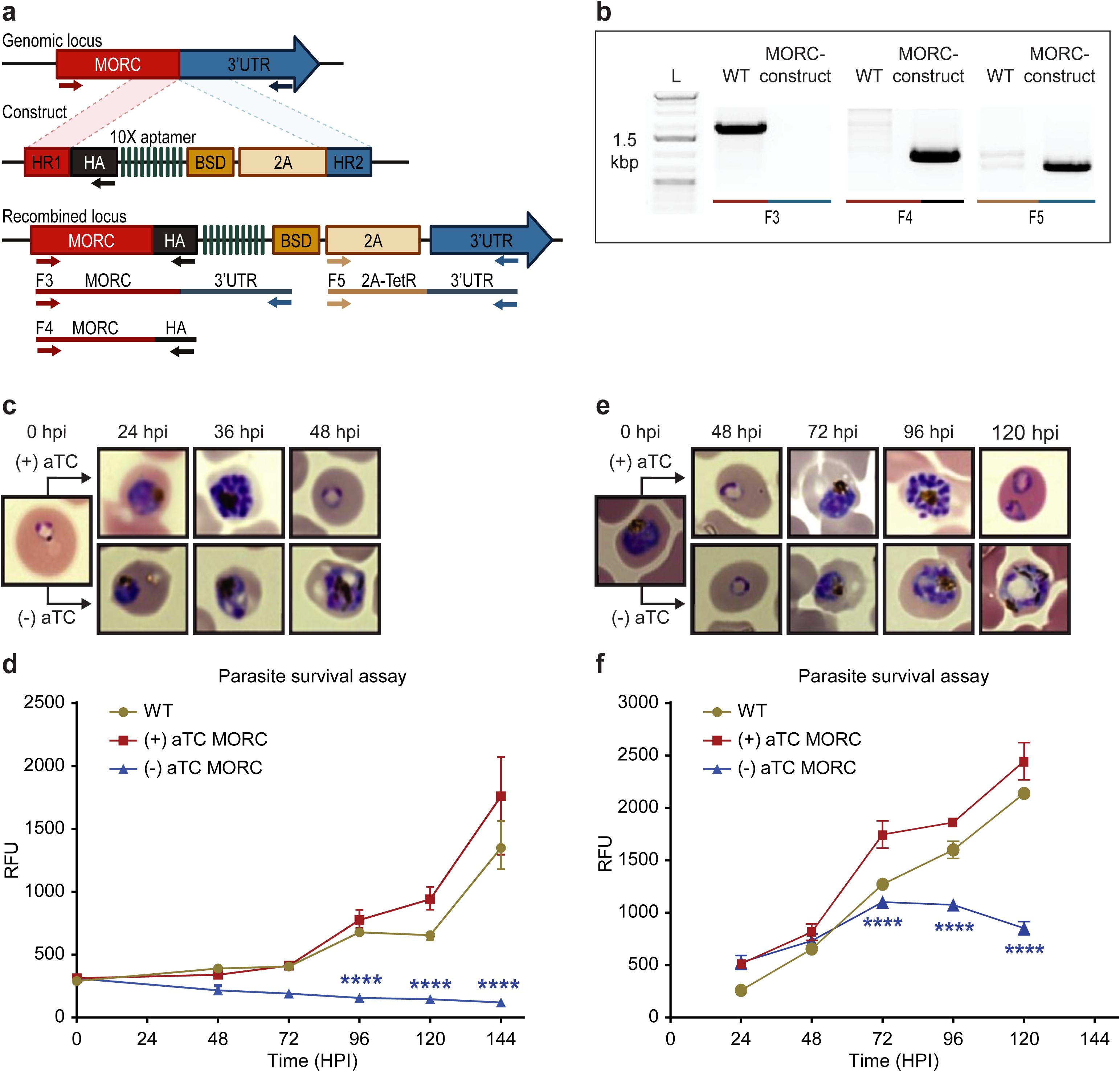
*Pf*MORC is essential for cell survival. **(a)** Diagram representation of *Pf*MORC-HA-TetR-DOZI plasmid. **(b).** PCR amplification is used to verify genomic insertion using primers sets targeting 1.5 kbp of WT *Pfmorc* genome locus absent in transgenic line (MORC construct) (F3) as well as verification of HA insertion (F4) and TetR-DOZI system extending along 3’ UTR of the construct (F5). **(c)** Phenotypic and **(d)** quantitative analysis of parasite cell progression after aTC withdrawal at the ring stage (0-6 hpi) (2-way ANOVA, n=3, p≤0.0001). **(e)** Phenotypic and quantitative **(f)** analysis of parasite cell progression after aTC removal at the trophozoite stage of cell cycle progression (24 hpi) (2-way ANOVA, n=3, p≤0.0001).

We then performed a phenotypic assay on (+/-) aTC *Pf*MORC-HA-TetR-DOZI parasite cultures. The assay was conducted in replicates of synchronized cultures that were then split into (+) aTC and (-) aTC conditions either at the ring or trophozoite stages on *Pf*MORC-HA-TetR-DOZI clones and WT control lines (**Fig. 3c-f, Sup. data 5**). Sequential morphological screens were performed through Giemsa-stained smears and monitored by microscope imaging. As opposed to what was observed previously with the glmS-ribozyme system(*41*), aTC removal at the ring stage induced clear signs of stress and cell cycle arrest in mid-trophozoite and schizont stages of the first intraerythrocytic cycle (**Fig. 3c-d**) compared to (+) aTC *Pf*MORC and WT parasites. Our data showed an ∼53%, 77%, and 84% (n=3, p≤0.0001) drop in parasitemia at 48-, 72- and 120-hours post invasion (hpi), respectively, when compared to control conditions. aTC supplemented and WT cultures showed unperturbed cell cycle progression, reinvasion, and morphological development. Interestingly, when aTC was withheld at the late trophozoite stages (24 to 30 hpi), parasites could complete the first cycle and successfully reinvade into new red blood cells (RBCs) (**Fig. 3e-f**). The evident hindered phenotypic response may either be associated with the delay time for the protein to be completely depleted or may suggest a more dispensable role of *Pf*MORC in the schizont stage. However, clear indications of stress and cell cycle arrest were ultimately detected at trophozoite and early schizont stages of the second cell cycle. Quantitative analysis of (-) aTC *Pf*MORC cultures revealed significantly decreased parasitemia of ∼37% and 66% (n=3, p≤0.0001) at 96 hpi and 120 hpi, respectively, compared to (+) aTC *Pf*MORC and WT control lines (**Fig. 3e-f, Sup. data 5**), confirming the importance of the protein at the trophozoite and early schizont stages.

### Effect of *Pf*MORC knockdown on parasite transcriptome

To define the effects of *Pf*MORC KD on transcription, we conducted detailed time-course measurements of mRNA levels using RNA-sequencing (RNA-seq) throughout the parasite asexual cycle on (+/-) aTC *Pf*MORC lines. Parasites were first synchronized and aTC was removed from one set of samples. Total RNA was then extracted at the trophozoite (24 hpi) and the schizont (36 hpi) stages to allow for detection of early and late changes in gene expression after aTC removal. Pairwise correlations analysis (**Fig. S4a**) between (+/-) aTC *Pf*MORC treated lines at the two different time points confirmed appropriate reproducibility of our RNA-seq experiments.

Our results identified a relatively low number of differentially expressed genes at 24 hpi with 96 and 93 upregulated and downregulated genes, respectively (FDR < 0.05 and log_2_ FC > 0.5 or < -0.5) (**Fig 4a, Sup. data 6**). This data is in agreement with the absence of major phenotypic changes observed in (-) aTC *Pf*MORC parasite cultures after 24 hours. GO enrichment analysis indicated upregulation of genes in response to xenobiotic stimulus including several phosphatases, hydrolases and heat shock proteins suggesting stress sensing of the parasites at this stage (**Fig. 4c**). GO analyses of downregulated genes, on the other hand, were found to be closely associated with regular metabolic processes and intracellular transport mechanisms suggesting dysregulation of regular cellular activity. Although, at this stage it is probable that the limited depletion of *Pf*MORC (58%) has restricted effects at the transcriptional level **(See Fig. S3)** and that the observed down-regulated genes are likely due to an indirect effect of cell cycle arrest. At 36 hpi the number of genes that exhibit changes in gene expression were significantly higher with 1319 upregulated and 1150 downregulated (FDR < 0.05 and log_2_ FC > 0.5 or < -0.5) in (-) aTC *Pf*MORC conditions, reflecting of a significant decrease of the protein ( ∼ 88.2%) and confirming the significant changes observed at the phenotypic level (**Fig. 4b**). These results emphasize the cascading effects of *Pf*MORC KD as the parasite progressed throughout its cell cycle. GO analysis indicated an enrichment of upregulated genes involved in multiple pathways including ATP metabolic process, mitochondria, translation, food vacuole and protein folding (**Fig. 4d**); characteristic of not only stress, but also a clear signal of cell cycle arrest in (-) aTC *Pf*MORC samples. Among upregulated genes, we also observed several *var* genes and genes exported at the surface of the red blood cell that could be linked to significant decreased in *Pf*MORC binding of these genes families as well as a disorganization of the heterochromatin cluster(s) at the trophozoite and schizont stages. GO enrichment analysis for genes that were downregulated included many genes involved in DNA replication, chromosome organization and mitotic spindle further emphasizing a strong cell cycle arrest and absence of potential major compensatory mechanisms for cell division and chromatin organization (**Fig. 4d**). Additional downregulated genes were required for invasion such as several merozoite surface proteins. Our results clearly indicate stress sensing and an arrest in cell cycle progression at the trophozoite and schizont stages (**Sup. data 5**)(*59*).

**Fig. 4.**
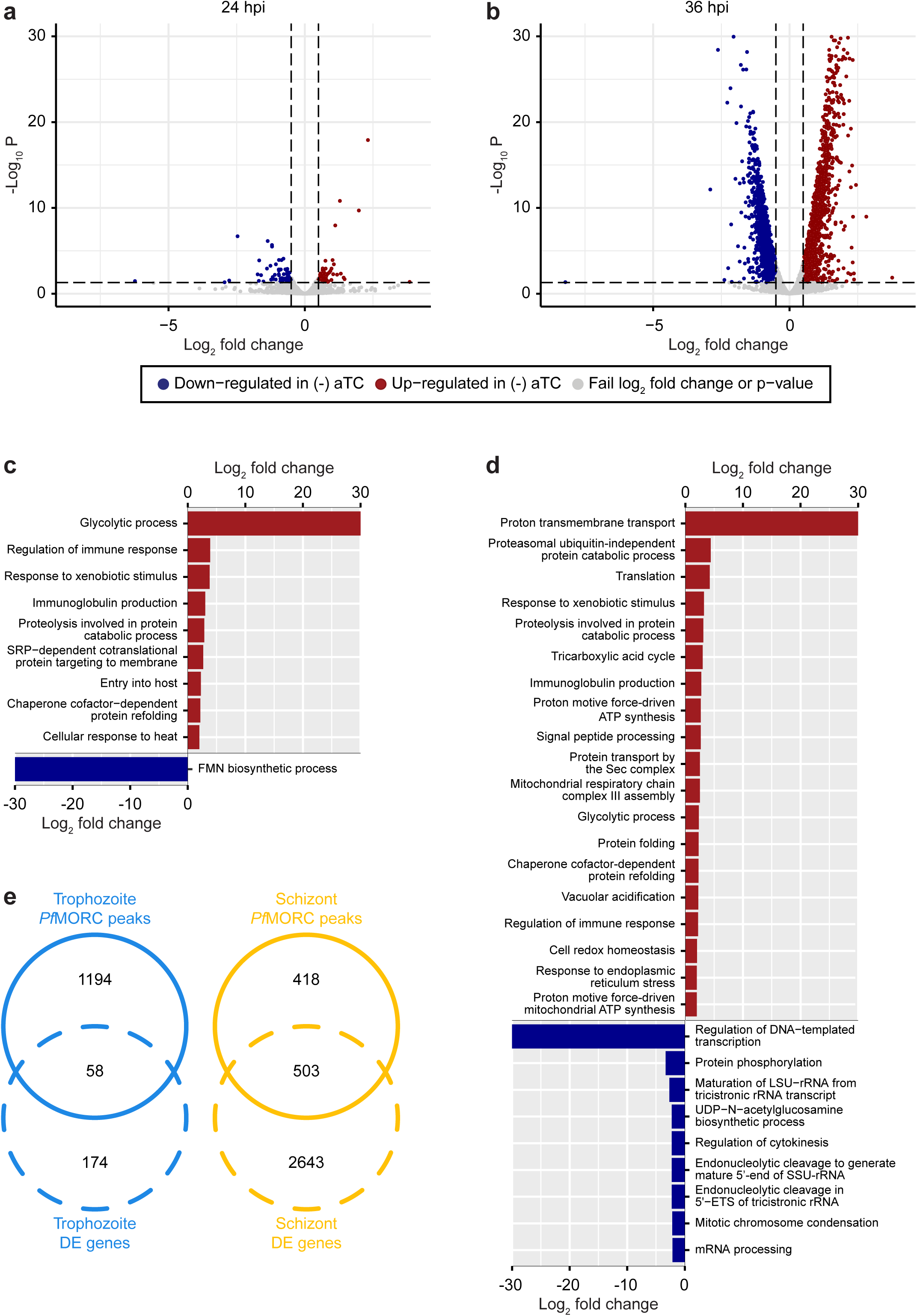
*Pf*MORC KD on parasite transcriptome. Volcano plots denoting upregulated (red), and downregulated (blue) genes discovered through differential expression analysis following *Pf*MORC knockdown at **(a)** 24 hpi **(b)** 36 hpi. Gene ontology enrichment analysis for upregulated (red) and downregulated (blue) genes at **(c)** 24 hpi and **(d)** 36 hpi. **(e)** Overlap of differentially expressed genes and genes containing significant peaks called by *Pf*MORC ChIP-seq analysis at the trophozoite and schizont stages.

Although these results are compelling, the small overlap observed between the ChIP-seq signals and RNA-seq results indicate that major changes observed in gene expression at the schizont stages are not simply the result of reduced *Pf*MORC binding in targeted gene bodies but a combination of direct and indirect effects of the degradation of *Pf*MORC that leads to cell cycle arrest and potential collapse of the heterochromatin (**Fig. 4e**).

### *Pf*MORC knockdown erodes antigenic-gene silencing framework

We next sought to analyze the effect of *Pf*MORC down-regulation on the global chromatin landscape. We therefore performed a ChIP-seq experiment against histone H3K9me3 and H3K9ac marks in response to *Pf*MORC depletion. Synchronized parasites were equally split between permissive and repressive conditions. At trophozoite (24 hpi) and schizont (36 hpi) stages of development, both control (+aTC) and experiment (-aTC) samples were fixed and parasites collected for chromatin immunoprecipitation followed by sequencing. The experimental procedure was performed in duplicate between (+/-) aTC *Pf*MORC treated lines. Correlation analysis confirms the reproducibility of our ChIP–seq experiments.

Results revealed no significant changes in histone H3K9ac marks across the genome (data not shown) but a reduction in the heterochromatin landscape in the *Pf*MORC depleted conditions, specifically in the telomere regions of the chromosomes **(Fig. 5a)**. *Pf*MORC KD resulted in significantly reduced (Mann-Whitney *U* test, p < 0.05) H3K9me3 marks within *var* gene promoters from 200-800 bp upstream of the TSS, as well as 400-800 bp downstream of the TSS **(Fig. 5b-c)**. This result coincides with our transcriptomic profile and the upregulation of *var* genes at 36 hpi of (-)aTC samples **(Fig 4d, Sup. data 6)**. Further analyses revealed similar trends in other gene families associated with parasite reinvasion and immune response typically silenced under WT conditions. Most noticeable was the reduced H3K9me3 coverage across the *rifin* gene family, which displayed a significant reduction (p < 0.05) in all bins from 1000 bp upstream of the TSS to 1000 bp downstream from the end of the gene **(Fig. 5b-c)** in response to *Pf*MORC depletion. There is a clear reduction of H3K9me3 coverage of virulence genes shown to be upregulated following *Pf*MORC knockdown **(Fig. 5c)**. Overall, these findings confirm that *Pf*MORC protein depletion not only affects chromatin landscape but has significant impact on heterochromatin. *Pf*MORC downregulation leads to the erosion of heterochromatin integrity most likely required for mutually exclusive gene expression and immune evasion within heterochromatin clusters.

**Fig. 5.**
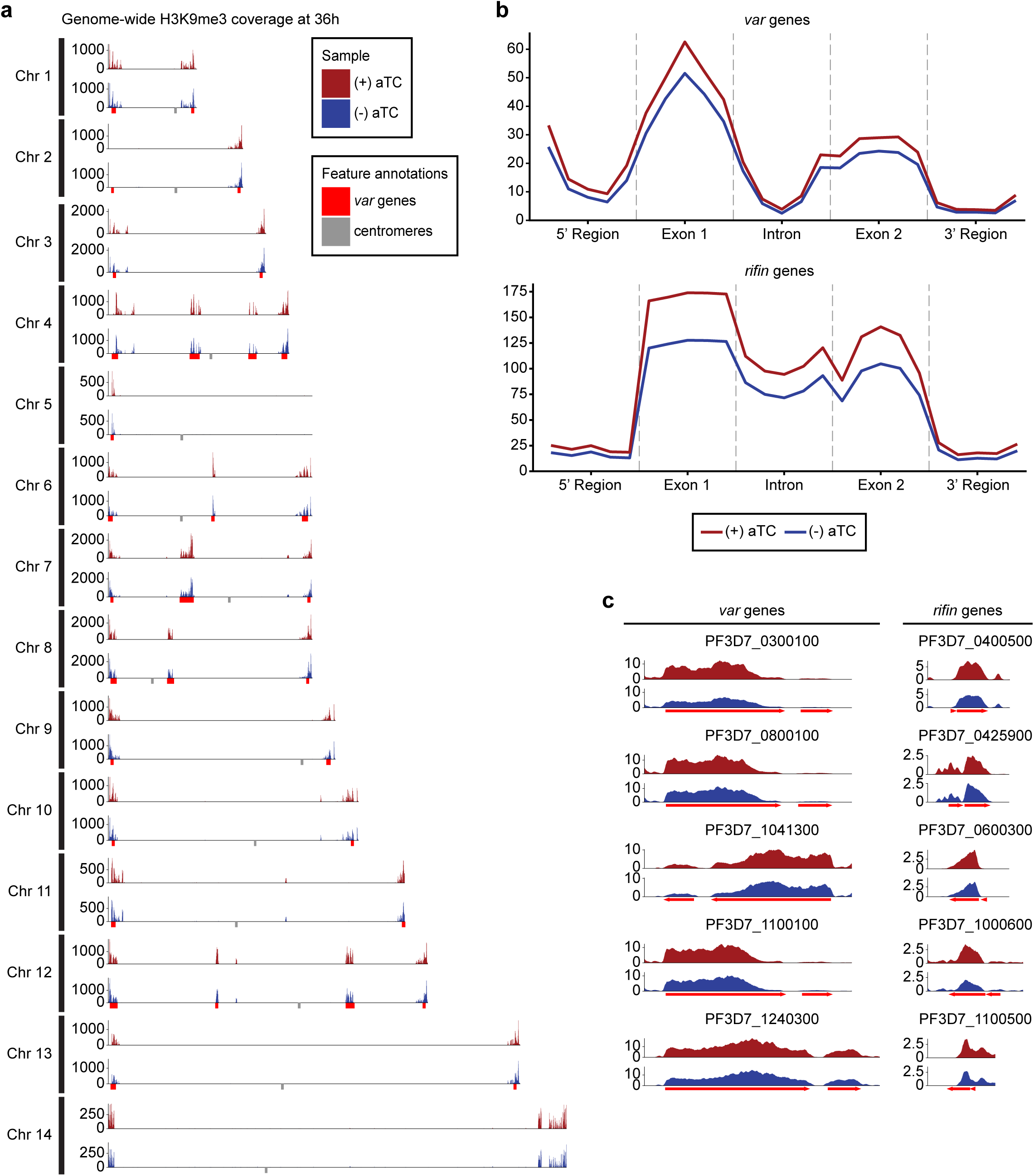
Impact of *Pf*MORC KD on heterochromatin markers. **(a)** Genome-wide H3K9me3 coverage of IGG subtracted and per-million normalized (+/-) aTC show similar distribution and concentration within telomeres and antigenic gene clusters highlighted in red. Replicates (n = 2) are merged using the mean of the normalized read coverage per base pair. **(b)** Binned coverage of *var* and *rifin* genes from 1 kb upstream of the TSS to 1 kb downstream of the end. The exons and intron of all genes within these families are split into five equal sized bins and the 5’ and 3’ regions are binned into five 200 bp bins. Read counts within each bin are per-million and bin length normalized prior to plotting. **(c)** Coverage of the five most upregulated *var* and *rifin* genes as determined by the transcriptomic analysis which shows elevated coverage in (+) aTC cells.

### Knockdown of *Pf*MORC expression results in the loss of tightly regulated heterochromatin structures

To better understand the effect that downregulation of *Pf*MORC has on the chromatin structure and investigate whether changes in chromatin accessibility may explain large changes in gene expression observed using RNA-seq, we performed Hi-C on (+/-) aTC *Pf*MORC cultures at 24 hpi (trophozoite) and 36 hpi (schizont). Biological replicates for each sample were collected and used to generate Hi-C libraries with >37 million reads per replicate/sample, ensuring comprehensive coverage of both intrachromosomal and interchromosomal interactions. After processing (pairing, mapping, and quality filtering) the raw sequences via HiC-Pro (*60*), there were approximately 15 million (σ = ∼10 million) high quality interaction pairs per sample. Due to the high number of reads and relatively small size of the *P. falciparum* genome (23.3 Mb) compared to higher eukaryotes, we elected to bin our Hi-C data at 10-kb resolution, which allowed the identification of genome-wide patterns while not introducing much noise by binning at too high of a resolution. A high stratum-adjusted correlation (**Fig. S4**), especially at the 24 hpi time point, suggested that the chromatin structure was consistent between biological replicates, therefore we chose to combine replicates for downstream analyses and visualization.

Because of variation in the number of valid interaction pairs between (+/-) aTC *Pf*MORC samples, 100 iterations of random sampling were performed on samples with higher read count to obtain ∼35 million and ∼9 million interaction pairs at 24 hpi and 36 hpi, respectively. ICED normalized intrachromosomal heatmaps displayed a high proportion of interactions at distances less than 10% the length of each chromosome as well as strong subtelomeric interactions and internal regions containing genes involved in antigenic variation (**Fig. 6a-b, Fig. S6-S9**) This pattern was similar to those observed previously at various stages of the life cycle (*27, 60, 61*) and confirm that genes involved in antigenic variation are usually confined within dense heterochromatin rich regions to aid the tight control of *var* genes necessary for their mutually exclusive expression and immune evasion(*53, 54*). Data generated at 36 hpi in (-) aTC *Pf*MORC parasite cultures were however tumultuous, with a decrease in defined heterochromatin borders and fewer long-range intrachromosomal interactions across the genome. This may indicate a significant loss of chromatin maintenance in (-) aTC *Pf*MORC parasites (**Fig. 6a-b**).

**Fig. 6.**
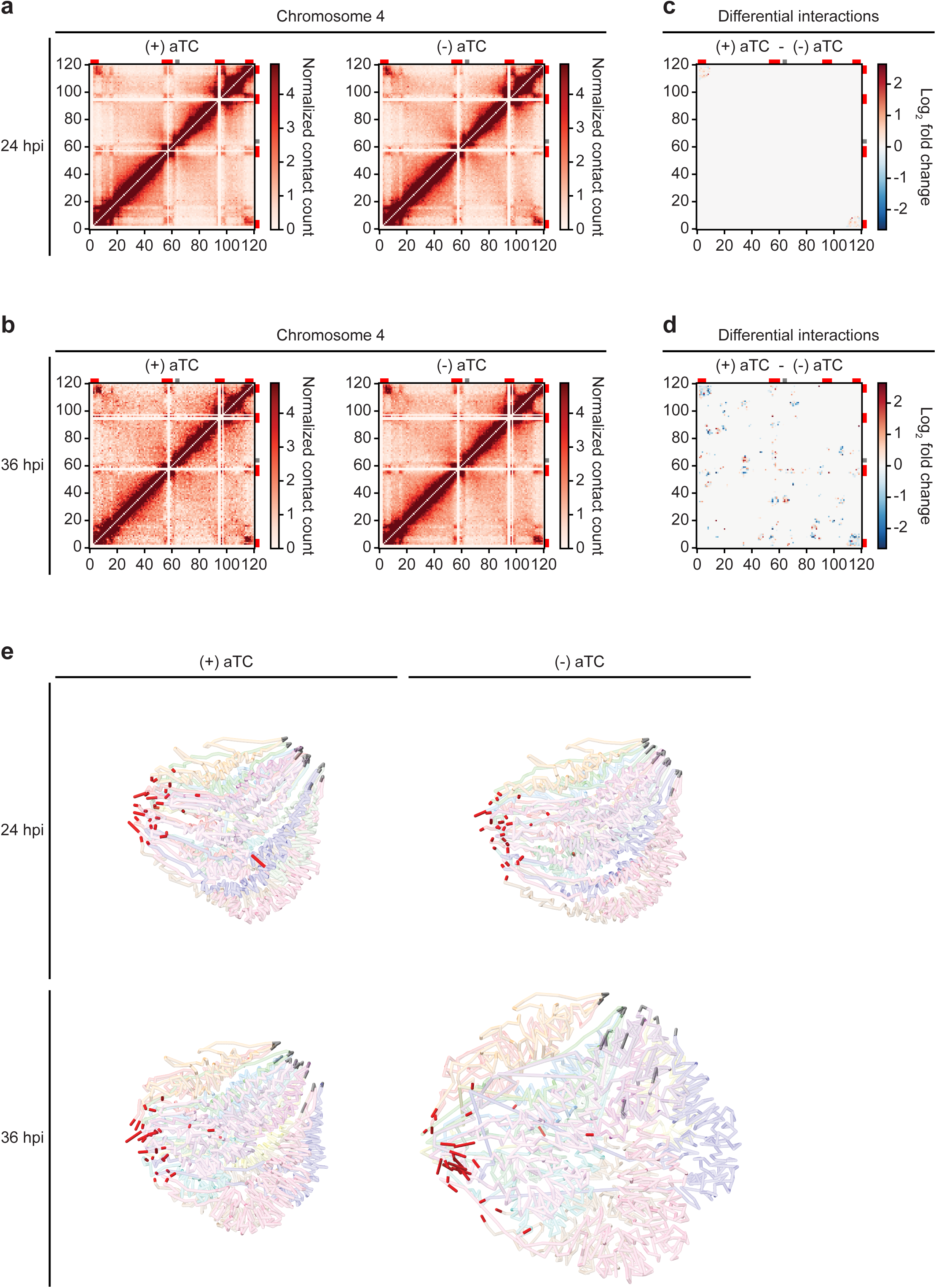
Loss of *Pf*MORC expression correlates with heterochromatin expansion. Intrachromosomal interaction heatmaps of (+/-) aTC *Pf*MORC for chromosome 4 at **(a)** 24 hpi and **(b)** 36 hpi displaying heterochromatin clustering within antigenic (*var*, *rifin* and *stevor*) gene-dense regions (red). Differential interaction heatmaps highlight changes in chromatin structure following removal of aTC and subsequent *Pf*MORC knockdown at **(c)** 24 hpi and **(d)** 36 hpi. **(e)** Whole-genome 3D models of the chromatin structure at both time points (24 hpi and 36 hpi) and (+/-) aTC.

To pinpoint which regions were most strongly affected by knockdown of *Pf*MORC, we used Selfish(*62*) to identify differential intrachromosomal and interchromosomal interactions in the (+/-) aTC *Pf*MORC samples. Although the 24 hpi and 36 hpi time points shared similarities in loci most highly affected by *Pf*MORC KD on many chromosomes, the changes at 36 hpi were more significant with a higher log_2_ FC (FDR < 0.05) at most loci (**Fig. 6c-d, Fig. S10-S11**). We also observed consistent loss of interactions between most regions containing *var* genes. These results indicate a failed attempt of the (-) aTC *Pf*MORC parasites to maintain their overall chromatin structure with a significant weakening of the tightly controlled heterochromatin cluster.

We further validated our results using PASTIS to generate coordinate matrices and subsequently visualize a consensus three-dimensional model of the chromosome folding and their overall organization within the nucleus (*63*). This can indicate changes in spatial distance that best describe the interaction data. The overall change in the chromatin 3D structure was clearly shown by our models of (+/-) aTC *Pf*MORC samples (**Fig. 6e**). While the co-localization of centromeres and telomeres in different regions of the nucleus were conserved, the 3D chromatin structure of (-) aTC *Pf*MORC at 36 hpi displayed clear opening of the chromatin and loss of interactions in most regions. This could explain the large changes in gene expression, including increased expression of all *var* and *rifin* genes in the (-) aTC *Pf*MORC line (**Sup. data 6**). While heterochromatin maintenance may be essential for tight control of *var* gene expression, preservation of the overall structure of the chromatin may be necessary to regulate the accurate expression of most *P. falciparum* transcripts.

## Discussion

Despite intensive investigations on the dynamic nature of the *Plasmodium* chromatin, progress has been slow in capturing the regulatory factors that maintain and control chromatin structure and gene expression throughout parasite development. Preliminary studies performed by our lab and others have identified *Pf*MORC as one of the most abundant chromatin-bound proteins(*32, 35*). MORC proteins from plant to animal cells have a broad range of binding sites within the genome and participate in either DNA methylation establishment or heterochromatin formation (*30, 64–69*). They have also been shown to co-localize with TFs in unmethylated promoter regions to alter chromatin accessibility and regulate TF binding and gene expression (*66–69*). For instance, AP2-P (PF3D7_1107800), AP2-G5, ISW1, NUP2, MCM5 and HDAC1 have been shown to interact with MORC in several published works (33, 42, 43), including this work. Protein pull-down experiments in *Plasmodium* spp. and *T. gondii* studies have detected MORC in complexes with several AP2 TFs. In *T. gondii, Tg*MORC was characterized as an upstream transcriptional repressor of sexual commitment (33, 35). In *Plasmodium*, the role of *Pf*MORC was still poorly understood. Conflicting results on the essentiality of *Pf*MORC using either KD or KO strategies remained unresolved(*32, 56*). Using the CRISPR/Cas9 genome editing tool, we have determined the nuclear localization, genome-wide distribution, and regulatory impacts of *Pf*MORC on the parasite chromatin, transcriptome, and cell cycle development. We now have powerful evidence demonstrating that *Pf*MORC is not only critical for parasite cell cycle progression and its survival but has a direct role in heterochromatin formation and gene silencing, including regulating immune evasion through antigenic variation. Disparities observed in the essentiality of *Pf*MORC for the parasite survival in previous studies(*32*) are most likely the results of weak protein disruption highlighting the need for a significant downregulation of *Pf*MORC for true functional analysis. Using IFA and protein pull-downs, we have confirmed that *Pf*MORC localizes to the nucleus and interacts with multiple ApiAP2 TFs, chromatin remodelers, and epigenetic players associated with heterochromatin regions including H3K9me3, SIP2, HDAC1 and the ISWI chromatin-remodeling complex (SWIB/MDM2)(*52*) (**Fig. 1**). Although some discrepancies have been observed with other studies, mainly due to the fact that MORC protein was not used as the bait protein, our study validated the interaction of MORC with these partners, supporting that *Pf*MORC is in complex with key heterochromatin regulators. Using a combination of ChIP-seq, protein knock down, RNA-seq and Hi-C experiments, we also demonstrated that the MORC protein is essential for the tight regulation of gene expression. We can speculate that lack of MORC impacts heterochromatin and chromatin compaction, preventing access to gene promoters from TFs and the general transcriptional machinery in a stage specific manner. Although additional experiments will be required, our hypothesis is reinforced by the fact that downregulation of the *Pf*MORC significantly reduced H3K9me3 coverage in the heterochromatin cluster(s) and 5’ flanking regions of *var* and *rifin* genes. While we were unable to confirm a direct role of *Pf*MORC in sexual conversion due to a nonsense mutation in AP2-G gene in our transfected lines, its strong interaction with AP2-G during the asexual cell cycle indicates that PfMORC in combination with other epigenetic factors may most likely control AP2-G expression and sexual differentiation. It is also important to recognize that the many pathways affected at the transcriptional level throughout the asexual stages in (-) aTC *Pf*MORC lines are not only the direct results of *Pf*MORC downregulation and decrease of its targeted DNA binding site. They are most likely a combination of direct and indirect effects of *Pf*MORC KD warranted by the cell cycle arrest observed at the phenotypic level and the collapse of the chromatin organization confirmed using chromatin conformation capture experiment. These direct and indirect effects should be carefully considered, and a combination of functional genomic studies should be completed when interpreting changes in gene expression in mutant generated lines.

All together our work demonstrated that in addition to its direct role in heterochromatin formation and antigenic variation, *Pf*MORC may act as a potential repressor to control a set of parasite specific genes including genes involved in parasite egress and invasion, and antigenic variation between the trophozoite to schizont stage transition(*52*). As our data confirm the importance of *Pf*MORC and the parasite specificity of several interacting partners, it is tempting to speculate that drugs targeting these protein complexes could lead to novel antiparasitic strategies.

## Methods

### Asexual parasite Culture and maintenance

Asexual *P. falciparum* strain NF54 or 3D7 parasites (MRA-1000, MRA-102 respectively) were propagated in 5% of human O^+^ erythrocytes and 10 mL of RPMI-1640 medium containing 0.5% Albumax II (Invitrogen), 2 mM L-glutamine, 50 mg/L hypoxanthine, 25 mM HEPES, 0.225% NaHCO_3_ and 10 mg/mL gentamicin. They were maintained at 37°C and gassed with a sterile mixture of 5% O_2_, 5% CO_2_ and 90% N_2_(*70, 71*). Parasite synchronization was achieved using two 5% D-sorbitol treatments 8 hours apart(*70, 71*).

### Plasmid Construction

Flagging of the *P. falciparum* MORC (Pf3D7_1468100) gene spanning position (Ch14: 2785386 - 2797934 (-)) was performed using a two-plasmid design to insert a 3x HA tag. The pCasG-Cas9-sgRNA plasmid vector (gifted by Dr. Sean Prigge) contains the site to express the sgRNA, along with the yDHODH gene as the positive selection marker. The gRNA oligos (**Sup. data 1**) were ligated after digestion of the plasmid with BsaI. The pDC2-cam-Cas9-U6 plasmid (gifted by Dr. Marcus Lee) was digested with BamHI and Apal to remove the eGFP tag from the backbone. 453 bp of C-terminal region of *Pf*MORC and 469 bp of 3’UTR were amplified from *P. falciparum* genomic DNA with their respective primers (**Sup. data 1**). A 3x HA-tag was fused to the C-terminal region of the amplified product along with the 3’UTR formed through Gibson assembly master mix (NEB, E2611S). To generate the *Pf*MORC-HA knockdown constructs, a pKD*^Pf^*^AUBL^ plasmid (gifted by Dr. Sean Prigge)(*72*) was digested with AscI and AatII. Homology arms HA1 and HA2 of *Pf*MORC were amplified with respective primers (**Sup. data 1**) having 20 bp overhang and inserted into the digested plasmid using Gibson assembly mix. The resulting pKD-MORC-HA-TetR-DOZI construct was linearized with EcoRV enzyme prior to transfection. All constructs were confirmed through restriction enzyme digestion and Sanger sequencing.

Plasmids were isolated from 250 mL cultures of *Escherichia coli* (XL10-Gold Ultracompetent Cells, Agilent Cat. 200314) and 60 µg of pDC2-MORC-HA or linearized pKD-MORC-HA-TetR-DOZI, were used with 60 µg of pCasG-plasmid containing gRNA to transfect 200 µl of fresh red blood cells (RBCs) infected with 3-5% early ring stage parasites. After one erythrocytic cycle, transfected cultures were supplemented with 1.5 µM WR99210 (2.6nM) (provided by the (Jacobus Pharmaceuticals, Princeton, NJ) and 2.5 μg/mL Blasticidin (RPI Corp B12150-0.1). *Pf*MORC-HA-TetR-DOZI transfected parasites were maintained with 500 nM anhydrotetracycline (aTC)(*45, 58*). Media and drug selection was replenished every 24 hours for 7 consecutive days after which DSM-1 drug selection was halted Once parasites were detected by microscopy, integration of the insert was confirmed by PCR amplification. To generate genetically homogenous parasite lines, the transfected parasites were serially diluted to approximately 0.5% parasite/well, into 96 well plates.

### Molecular analysis of the transgenic lines

Genomic DNA (gDNA) was extracted and purified using the DNeasy Blood & Tissue kit (Qiagen) following instructions from the manufacturer. The diagnostic PCR analysis was used to genotype the transfected lines using the primers listed in Supplementary data 1. The PCR amplification was conducted using KAPA HiFi HotStart ReadyMix (Roche) and amplicons were analyzed by gel electrophoresis followed by sequencing. For whole genome sequencing, genomic DNA was fragmented using a Covaris S220 ultrasonicator and libraries were generated using KAPA LTP Library Preparation Kit (Roche, KK8230). To verify that the insertion was present in the genome at the correct location in both transfected lines, reads were mapped using Bowtie2 (v2.4.4) to the *P. falciparum* 3D7 reference genome (PlasmoDB, v48), edited to include the insertion sequence in the intended location. Integrative Genomic Viewer (IGV, Broad Institute) was used to verify that reads aligned to the modified sequence.

### Variant analysis by genome-wide sequencing

To call variants (SNPs/indels) in the transfected lines compared to a previously sequenced control 3D7 line, genomic DNA reads were first trimmed to adapters and aligned to the *Homo sapiens* genome (assembly GRCh38) to remove human-mapped reads. Remaining reads were aligned to the *P. falciparum* 3D7 genome using bwa (version 0.7.17) and PCR duplicates were removed using PicardTools (Broad Institute). GATK HaplotypeCaller (https://gatk.broadinstitute.org/hc/en-us) was used to call variants between the sample and the 3D7 reference genome for both the transfected lines and the NF54 control. Only variants that were present in both transfected lines but not the NF54 control line were kept. We examined only coding-region variants and removed those that were synonymous variants or were located in *var*, *rifin*, or *stevor* genes. Quality control of variants was done by hard filtering using GATK guidelines.

### Immunofluorescence assays

A double-staining immunofluorescence assay was used on NF54 control and transgenic parasite lines of mixed parasite population growth stages. Parasites were washed in incomplete medium prior to fixing onto coverslips with 4% paraformaldehyde for 20 min at RT under darkness. After fixation, samples were washed 3-5 times with 1x PBS followed by permeabilization with 0.5%Triton X-100 in PBS for 25 min at RT. Subsequently, samples were subjected to PBS washes and then incubated overnight at 4°C in blocking buffer (2mg/ml BSA solution in PBS containing 0.05% Tween-20 solution). Following overnight blocking, samples were washed and incubated for 1 hr at RT with Anti-HA Rb Ab (1:500, Abcam, ab9110) in blocking buffer. After primary Ab incubation, samples were subject to 3x washes with wash buffer (1x PBS containing 0.05% Tween-20) followed by incubation with anti-rabbit DyLight 550 (Abcam ab98489; 1: 500) secondary antibody for 1 h at room temperature. After incubation and series of washes with wash buffer, slides were incubated with anti-H3K9me3 antibody, Alexa Fluor 488 conjugate (Millipore 07–442-AF488; 1:100) for 1 hr at RT. Slides are then washed and mounted in Vectashield Antifade Mounting Medium with DAPI (Vector Laboratories, H-1200). Images were acquired using a Keyence BZ-X810 Fluorescence Microscope and were processed through ImageJ.

### Western blotting

Parasites were synchronized twice at 8 hr. intervals between synchronizations. After one cycle, culture was washed 3x with complete media to remove residual aTC followed by dividing the cultures into two equal conditions. Parasites were grown with or without 500 nM aTC for 24 and 36 hrs. after which RBCs were lysed using 0.15 % saponin and parasites were collected after being washed with ice cold 1X PBS. Proteins were recovered from the lysed parasites after 30 min of incubation in lysis buffer (150 mM NaCl, 0.5 % NP40, 50 mM Tris pH 8, 1 mM EDTA and protease inhibitors) and 10 sec of sonication. Proteins were quantified with Pierce™ BCA Protein Assay Kit (Thermo Fisher 23227). 20 µg of proteins were loaded onto 3–8% Criterion XT Tris-Acetate Midi Protein Gels (Bio-Rad, 3450129). After migration, proteins were transferred onto a PVDF membrane and the membrane was blocked and then probed overnight with anti-HA tag antibody (1:2500, Abcam, ab9110) as well as anti-aldolase antibody as a loading control (1:10000, abcam, ab252953). After primary Ab incubation, blots were subsequently washed 3x with a washing buffer followed by HRP-labeled Goat anti-Rabbit IgG (H + L) (1:10,000, Novex^TM^, A16104). Clarity^TM^ Western ECL Substrate (Bio-Rad, 1705060) was applied to reveal the blots. Relative abundance of (+/-) aTC *Pf*MORC was calculated by Bio-Rad ChemiDoc imagelab software.

### Immunoprecipitation followed by MudPIT mass spectrometry

Mid-to late-stage asexual parasites were collected following saponin treatment and purified samples were then resuspended into fresh IP buffer (50 mM Tris-HCl pH 7.5, 300 mM NaCl, 0.5 mM EDTA, 0.5 mM EGTA, 2 mM AEBSF 0.5 % Triton X-100, and EDTA-free protease inhibitor cocktail (Roche)). Post cell lysis solution was homogenized via sonication for 6-9 rounds. The soluble extracts were centrifuged at 13,000 x g for 15 min at 4°C. The lysates were precleared with Dynabeads™ Protein A (Invitrogen) for 1h at 4°C. Anti-HA tag antibody (1:2500, Abcam, ab9110) are added to control and HA tagged *Pf*MORC precleared protein extract samples for 1hr at 4°C under constant rotation followed by the addition of fresh Dynabeads™ Protein A beads to each sample and incubated overnight at 4°C. Dynabeads were washed 3 times with 500 µL of buffer (PBS, 0.05% Tween-20). Proteins were eluted into an elution buffer (50 mM Tris-HCl pH 6.7, 100 mM DTT and 2% SDS). The eluent was subsequently precipitated overnight in 20% TCA followed by cold acetone washes. The urea-denatured, reduced, alkylated, and digested proteins were analyzed by Multidimensional Protein Identification Technology (MudPIT) on a Orbitrap Elite mass spectrometer coupled to an Agilent 1260 series HPLC, as described previously (*73*).

### Proteomics data processing and analysis

Tandem mass (MS/MS) spectra were interpreted using ProluCID v.1.3.3 (*74*) against a database consisting of 5527 non-redundant (NR) Plasmodium falciparum 3D7 proteins (PlasmoDB, v42), 36661 NR human proteins (NCBI, 2018-03-30 release), 419 common contaminants (human keratins, IgGs, and proteolytic enzymes), together with shuffled versions of all of these sequences. DTASelect v.1.9 (*75*) and swallow v.0.0.1, an in-house developed software (https://github.com/tzw-wen/kite) were used to control FDRs resulting in protein FDRs less than 1.86%. All datasets were contrasted against their merged data set, respectively, using Contrast v1.9 (*75*) and in-house developed sandmartin v.0.0.1 (https://github.com/tzw-wen/kite/tree/master/ kitelinux). Our in-house developed software, NSAF7 v.0.0.1 (https://github.com/tzw-wen/kite/tree/master/windowsapp/NSAF7x64), was used to generate spectral count-based label free quantitation results (*76*). QPROT(*49, 77*) was used to calculate values of log_2_ fold change and Z-statistic to compare two replicate *Pf*MORC affinity purifications to two negative controls. Proteins enriched in the *Pf*MORC-HA samples with values of log_2_ fold change ≥ 2 and Z-statistic ≥ 5 were considered significantly enriched.

### ChIP assay

*Pf*MORC-HA and *Pf*MORC KD asexual stage parasites were supplemented with and without aTC along with NF54 parasites (as a control) and were harvested at 24 and 36 HPS and cross linked with 1% formaldehyde for 10 min at 37°C followed by quenching with 150 mM glycine and 3 x washing with 1 x PBS. The pellets were resuspended in 1 mL of nuclear extraction buffer (10 mM HEPES, 10 mM KCl, 0.1 mM EDTA, 0.1 mM EGTA, 1 mM DTT, 0.5 mM AEBSF and 1x Protease inhibitor cocktail), and incubated on ice for 30 min before addition of NP-40/Igepal to a final of 0.25%. After lysis, the parasites are subject to homogenization by passing through a 26G 3/8 needle/syringe to burst the nucleus. After centrifugation for 20 min at 2,500 x g at 4°C, samples were resuspended in 130 µL of shearing buffer (0.1% SDS, 1 mM EDTA, 10 mM Tris-HCl pH 7.5 and 1X Protease inhibitor cocktail) and transferred to a 130 µL Covaris sonication microtube. The samples were then sonicated using a Covaris S220 Ultrasonicator for 8 min (Duty cycle: 5%, intensity peak power:140, cycles per burst: 200, bath temperature: 6°C). The samples were transferred to ChIP dilution buffer (30 mM Tris-HCl pH 8.0, 3 mM EDTA, 0.1% SDS, 30 mM NaCl, 1.8% Triton X-100, 1X protease inhibitor tablet, 1X phosphatase inhibitor tablet) and centrifuged for 10 min at 13,000 rpm at 4°C, retaining the supernatant. For each sample, 13 μL of protein A agarose/salmon sperm DNA beads were washed three times with 500 µL ChIP dilution buffer by centrifuging for 1 min at 1,000 rpm at room temperature. For pre-clearing, the diluted chromatin samples were added to the beads and incubated for 1 hour at 4°C with rotation, then pelleted by centrifugation for 1 min at 1,000 rpm. Before adding antibody, ∼10% of each sample was taken as input. 2 µg of anti-HA tag (1:2500, Abcam, ab9110) or rabbit polyclonal anti-H3K9me3 or anti-H3K9ac (Millipore no. 07–442 and H0913) antibodies were added to the sample and incubated overnight at 4°C with rotation. For each sample, rabbit IgG antibody (Cat.689 No. 10500c, Invitrogen) was used as a negative-control library. Per sample, 25 µL of protein A agarose/salmon sperm DNA beads were washed with ChIP dilution buffer (no inhibitors), blocked with 1 mg/mL BSA for 1 hour at 4°C, and then washed three more times with buffer. 25 µL of washed and blocked beads were added to the sample and incubated for 1 hour at 4°C with continuous mixing to collect the antibody/protein complex. Beads were pelleted by centrifugation for 1 min at 1,000 rpm at 4°C. The bead/antibody/protein complex was then washed with rotation using 1 mL of each buffer twice; low salt immune complex wash buffer (1% SDS, 1% Triton X-100, 2 mM EDTA, 20 mM Tris-HCl pH 8.0, 150 mM NaCl), high salt immune complex wash buffer (1% SDS, 1% Triton X-100, 2 mM EDTA, 20 mM Tris-HCl pH 8.0, 500 mM NaCl), and TE wash buffer (10 mM Tris-HCl pH 8.0, 1 mM EDTA). Complexes were eluted from antibodies by adding 250 μL of freshly prepared elution buffer (1% SDS, 0.1 M sodium bicarbonate). 5 M NaCl were added to the eluate and cross-linking was reversed by heating at 45°C overnight followed by addition of 15 μL of 20 mg/mL RNase A with 30 min incubation at 37°C. After this, 10 μL of 0.5 M EDTA, 20 μL of 1 M Tris-HCl pH 7.5, and 2 μL of 20 mg/mL proteinase K were added to the eluate and incubated for 2 hours at 45°C. DNA was recovered by phenol/chloroform extraction and ethanol precipitation. DNA was purified using Agencourt AMPure XP beads. Libraries were then prepared from this DNA using KAPA LTP Library Preparation Kit (Roche, KK8230) and sequenced on a NovaSeq 6000 machine.

### ChIP-seq analysis

FastQC (version 0.11.8, https://www.bioinformatics.babraham.ac.uk/projects/fastqc/) was used to analyze raw read quality. Any adapter sequences were removed using Trimmomatic (version 0.39, http://www.usadellab.org/cms/?page=trimmomatic). Bases with Phred quality scores below 20 were trimmed using Sickle (version 1.33, https://asexualstageparasitesb.com/najoshi/sickle). The resulting reads were mapped against the *P. falciparum* genome (v48) using Bowtie2 (version 2.4.4). Using Samtools (version 1.11), only properly paired reads with mapping quality 40 or higher were retained and reads marked as PCR duplicates were removed by PicardTools MarkDuplicates (version 2.18.0, Broad Institute). Genome-wide read counts per nucleotide were normalized by dividing millions of mapped reads for each sample (for all samples including input) and subtracting IgG read counts from the anti-HA IP counts. Peak calling was performed using MACS3 (version 3.0.0a7) (*78*) with the upper and lower limits for fold enrichment set to 2 and 50, respectively.

### Phenotypic analyses

*P. falciparum* NF54 line along with transgenic *Pf*MORC-HA-TetR-DOZI line were synchronized, grown for one cycle post-synchronization and subject to 3 consecutive washes with 1X PBS before being supplemented either with or without (aTC) for 48 hrs. or 72 hrs. (depending on which stage the parasites were washed). Culture media were replaced daily with or without aTC supplementation.

### Quantitative Growth assay

Synchronous cultures either ring or trophozoite stage at 0.5 % parasitemia were grown with or without aTC (500 nM) into a 96 well plate in triplicates for each time point. Parasites were collected at every 24 hrs. time for 3 cycles and subjected to SYBR green assay (Thermo Fisher, S7523).

### RNA-seq library preparation

Parasites at ring, trophozoite or schizont stage were extracted following saponin treatment before flash freezing. Two independent biological replicates were generated for each time point, culture condition and line. Total RNA was extracted with TRIzol*®* LS Reagent (Invitrogen) followed by incubation for 1 hr with 4 units of DNase I (NEB) at 37°C. RNA samples were visualized by RNA electrophoresis and quantified on Synergy™ HT (BioTek). mRNA was then purified using NEBNext® Poly(A) mRNA Magnetic Isolation Module (NEB, E7490) according to the manufacturer’s instructions. Libraries were prepared using NEBNext® Ultra™ Directional RNA Library Prep Kit (NEB, E7420) and amplified by PCR with KAPA HiFi HotStart Ready Mix (Roche). PCR conditions consisted of 15 min at 37°C followed by 12 cycles of 98°C for 30 sec, 55°C for 10 s and 62°C for 1 min 15 sec, then finally one cycle for 5 min at 62°C. The quantity and quality of the final libraries were assessed using a Bioanalyzer (Agilent Technology Inc). Libraries were sequenced using a NovaSeq 6000 DNA sequencer (Illumina), producing paired-end 100-bp reads.

### RNA-seq data processing and differential expression analysis

FastQC [https://www.bioinformatics.babraham.ac.uk/projects/fastqc/] was used to analyze raw read quality and thus 11 bp of each read and any adapter sequences were removed using Trimmomatic (v0.39) [http://www.usadellab.org/cms/?page=trimmomatic]. Bases were trimmed from reads using Sickle with a Phred quality threshold of 25 (v1.33) [https://github.com/najoshi/sickle] and reads shorter than 18 bp were removed. The resulting reads were mapped against the *P. falciparum* 3D7 genome (PlasmoDB, v53) using HISAT2 (v2-2.2.1) with default parameters. Uniquely mapped, properly paired reads with mapping quality of 40 or higher were retained using SAMtools (v1.11) [http://samtools.sourceforge.net/]. Raw read counts were determined for each gene in the *P. falciparum* genome using BedTools [https://bedtools.readthedocs.io/en/latest/<u>#</u>] to intersect the aligned reads with the genome annotation. Differential expression analysis was performed using DESeq2 to call up- and down-regulated genes (FDR < 0.05 and log_2_ FC > 0.5). Volcano plots were made using the R package EnhancedVolcano.

### Hi-C library preparation

(+/-) aTC parasites at 24 hpi and 36 hpi were cross linked using 1.25% formaldehyde for 15 minutes at 37°C in 10 mL total volume followed by quenching with 150 mM glycine. Parasites were then washed three times with chilled 1x PBS on a rocking platform. Parasite nuclei were released using lysis buffer (10 mM Tris-HCl, pH 8.0, 10 mM NaCl, 2 mM AEBSF, 0.25% Igepal CA-630, and 1X EDTA-free protease inhibitor cocktail (Roche)) and 15 syringe passages through a 26.5-gauge needle. After releasing the crosslinked chromatin with 0.1% SDS, chromatin was digested using MboI restriction enzyme overnight at 37°C. Thereafter Hi-C libraries were prepared as previously described (*27, 79*).

### Hi-C data processing, differential interaction analysis and generation of 3D models

Paired-end Hi-C libraries were processed (pairing, mapping, quality filtering, binning, and normalization) using HiC-Pro(*60*). A mapping quality cutoff of 30 was set while aligning to the *P. falciparum* genome (PlasmoDB, v58) and the resulting reads were binned at 10 kb resolution and ICED-normalized. Stratum-adjusted correlation coefficients were calculated using HiCRep (*80*), then replicates were merged to generate a single consensus sample for each time point and condition. The normalized matrices were per-million read count normalized and maximum values for the heatmap color scale on each chromosome was set to the maximum value for all samples to allow for direct comparisons between each condition and time point. Furthermore, due to the overrepresentation of short-range interactions, the maximum values for each heatmap were also set to the 90th percentile of each chromosome to slightly compress highly interacting regions and enhance visualization of interactions. Significant intrachromosomal interactions were identified with FitHiC(*81*) to identify the log-linear relationship between contact probability and genomic distance. Differential interchromosomal and intrachromosomal interactions between (+) aTC and (-) aTC were identified using Selfish (*62*) with default parameters (FDR < 0.05). The maximum and minimum values for the color scales in the differential heatmaps were set to the absolute value of the largest log_2_ fold change within each chromosome. PASTIS (*63*) was used to generate coordinate matrices from the raw read count matrices output by HiC-Pro, and then visualized as 3D chromatin models in ChimeraX (*82*), highlighting regions containing *var* genes, telomeres, and centromeres.

### Statistical analyses

Parasitemia and proportion of asexual stages were analyzed using a two-way ANOVA with Tukey’s test for multiple comparisons. Significant differences were indicated as following: * for p < 0.05; ** for p < 0.01, *** for p < 0.001 and **** for p < 0.0001. Statistical tests were performed with GraphPad Prism. Figures were generated with GraphPad Prism and BioRender.

## Data Availability

Sequence reads for all sequencing experiments have been deposited in the NCBI Sequence Read Archive with accession PRJNA994684. Original data underlying this manuscript generated at the Stowers Institute can be accessed from the Stowers Original Data Repository at http://www.stowers.org/research/publications/LIBPB-2404. The mass spectrometry dataset generated for this study is available from the MassIVE data repository (ftp://MSV000092353@massive.ucsd.edu) using the identifier MSV000092353. All other data are available in the main text, methods, and supplementary data.

## Acknowledgement

This work was supported by NIH grants to KGLR (nos. 1R01 AI136511 and R21 AI142506-01) and by the University of California, Riverside to KGLR (no. NIFA-Hatch-225935). This publication includes data generated at the UC San Diego IGM Genomics Center utilizing an Illumina NovaSeq 6000 that was purchased with funding from a National Institutes of Health SIG grant (#S10 OD026929).

## Author contributions

Conceptualization was the responsibility of KGLR who also supervised the project together with LF. MG generated all the constructs and cell lines used in the study. Immunofluorescence and growth assays were performed by MG and ZC. ChIP seq, RNA seq libraries were generated by MG and analyzed by TL and SA. Hi-C experiments were performed by MG and the data was analyzed by TL. IP-MS experiments were performed by MG, ZC, CB, and analyzed by TH, CB and LF. Software and data curation were provided by TL, SA, and CB. ZC, TH, TL and KGLR wrote the original draft. All authors reviewed and edited the final version of the manuscript.

## Supplementary Data

**Supplementary data 1. *Pf*MORC plasmid constructs to generate transgenic lines.** All constructs were performed through listed primers, restriction sites and gRNAs.

**Supplementary data 2. Mapped Whole Genome sequencing results of *Pf*MORC transfectants.**

**Supplementary data 3. (a) Proteins identified by MudPIT analysis after MORC-HA immunoprecipitation.** *Pf*MORC (PF3D7_1468100) is highlighted in yellow and significantly purified proteins are in red. The filters used are QPROT log_2_ fold change > 2 and Z statistic > 5. Detailed protein list is provided in an additional sheet. **(b)** Raw data showing proteins identified by MudPIT analysis after *Pf*MORC-HA immunoprecipitation.

**Supplementary data 4. Peak calling result of *Pf*MORC ChIP-seq.** Results obtained at **(a)** ring stages **(b)** trophozoite stages and **(c)** schizont stages of cell progression.

**Supplementary data 5. Parasite Survival Assay illustrating effect of *Pf*MORC down-regulation** on parasites at ring stage (top) or trophozoite stage (bottom) cell cycle. Experiments were conducted in triplicates with parasitemia quantified via Relative Fluorescence Units (RFUs) obtained by SYBR green assay for each time point. Significance derived through 2-way ANOVA.

**Supplementary data 6. (a) RNA-seq read counts of *Pf*MORC at 24 and 36 hours.** DEseq2 analysis of differentially expressed genes of *Pf*MORC at **(b)** 24 hours and **(c)** 36 hrs.

**Figure S1. Expression of *Pf*MORC-HA in WT control and *Pf*MORC-HA lines.** Western blot image revealed expression of *Pf*MORC-HA protein in cultures using anti-HA antibody.

**Figure S2. Correlation, peak calling, and gene family specific coverage of *Pf*MORC ChIP- seq. (a)** Heatmap indicating correlation between time points and replicates. **(b)** *Pf*MORC coverage of select *var* genes from 500 bp 5’ of the TSS to 500 bp 3’ of the coding region. **(c)** *Pf*MORC coverage of three highly expressed *msp* genes from 500 bp 5’ of the TSS to 500 bp 3’ of the coding region.

**Figure S3. Expression of *Pf*MORC-HA in *Pf*MORC-HA-TetR-DOZI lines.** Western blot image reveals expression of *Pf*MORC-HA (Top) or WT protein (Bottom) in samples extracted at 24 HPS or 36 HPS in +/- aTC conditions using anti-HA antibody. WT samples loaded with exactly 50% of the total protein obtained in *Pf*MORC samples at each condition.

**Figure S4. RNA-seq correlation heatmap. (a)** Heatmap indicating correlation between (+/-) aTC condition, time points and replicates.

**Figure S5. Hi-C correlation analyses. (a)** Stratum-adjusted correlation between (+/-) aTC condition, time points and replicates. **(b)** Negative log-linear relationship between contact probability and genomic distance.

**Figure S6-9. Hi-C interaction heatmaps binned at 10kb resolution.** Each sample includes intrachromosomal contact count heatmaps for all 14 chromosomes within the *P. falciparum* genome and a genome-wide interchromosomal interaction heatmap. Data for each heatmap is ICED and per-million read count normalized, and the maximum y-value for each heatmap is set to the highest value within the given chromosome for all samples. Antigenic gene-containing regions (red) and centromeres (gray) are highlighted.

**Figure S10-S11. Differential Hi-C interaction heatmaps binned at 10kb resolution.** Heatmaps generated from differential interaction matrices, identifying regions with a positive (red) or negative (blue) log_2_ fold change between (-) aTC and (+) aTC. Data for each heatmap is ICED and per-million read count normalized, and the maximum y-value for each heatmap is set to the highest value within the given chromosome for all samples. Antigenic gene-containing regions (red) and centromeres (gray) are highlighted.

## References

1. WHO, World malaria report 2023. (2023).

2. R. M. Coulson, N. Hall, C. A. Ouzounis, Comparative genomics of transcriptional control in the human malaria parasite Plasmodium falciparum. Genome Res 14, 1548–1554 (2004).

3. S. Balaji, M. M. Babu, L. M. Iyer, L. Aravind, Discovery of the principal specific transcription factors of Apicomplexa and their implication for the evolution of the AP2-integrase DNA binding domains. Nucleic Acids Res 33, 3994–4006 (2005).

4. E. K. De Silva et al., Specific DNA-binding by apicomplexan AP2 transcription factors. Proc Natl Acad Sci U S A 105, 8393–8398 (2008).

5. M. Yuda et al., Identification of a transcription factor in the mosquito-invasive stage of malaria parasites. Mol Microbiol 71, 1402–1414 (2009).

6. S. Iwanaga, I. Kaneko, T. Kato, M. Yuda, Identification of an AP2-family protein that is critical for malaria liver stage development. PLoS One 7, e47557 (2012).

7. A. Sinha et al., A cascade of DNA-binding proteins for sexual commitment and development in Plasmodium. Nature 507, 253–257 (2014).

8. B. F. Kafsack et al., A transcriptional switch underlies commitment to sexual development in malaria parasites. Nature 507, 248–252 (2014).

9. K. M. Lesage et al., Cooperative binding of ApiAP2 transcription factors is crucial for the expression of virulence genes in Toxoplasma gondii. Nucleic Acids Res 46, 6057–6068 (2018).

10. M. Yuda, S. Iwanaga, S. Shigenobu, T. Kato, I. Kaneko, Transcription factor AP2-Sp and its target genes in malarial sporozoites. Mol Microbiol 75, 854–863 (2010).

11. K. Modrzynska et al., A Knockout Screen of ApiAP2 Genes Reveals Networks of Interacting Transcriptional Regulators Controlling the Plasmodium Life Cycle. Cell Host Microbe 21, 11–22 (2017).

12. C. Gu et al., Multiple regulatory roles of AP2/ERF transcription factor in angiosperm. Bot Stud 58, 6 (2017).

13. M. Yuda, I. Kaneko, S. Iwanaga, Y. Murata, T. Kato, Female-specific gene regulation in malaria parasites by an AP2-family transcription factor. Mol Microbiol 113, 40–51 (2020).

14. M. Yuda, S. Iwanaga, I. Kaneko, T. Kato, Global transcriptional repression: An initial and essential step for Plasmodium sexual development. Proc Natl Acad Sci U S A 112, 12824–12829 (2015).

15. Z. Bozdech et al., Expression profiling of the schizont and trophozoite stages of Plasmodium falciparum with a long-oligonucleotide microarray. Genome Biol 4, R9 (2003).

16. K. G. Le Roch et al., Discovery of gene function by expression profiling of the malaria parasite life cycle. Science 301, 1503–1508 (2003).

17. Z. Bozdech et al., The transcriptome of the intraerythrocytic developmental cycle of Plasmodium falciparum. PLoS Biol 1, E5 (2003).

18. P. Srinivasan et al., Analysis of the Plasmodium and Anopheles transcriptomes during oocyst differentiation. J Biol Chem 279, 5581–5587 (2004).

19. F. Silvestrini et al., Genome-wide identification of genes upregulated at the onset of gametocytogenesis in Plasmodium falciparum. Mol Biochem Parasitol 143, 100–110 (2005).

20. M. Yang et al., Full-Length Transcriptome Analysis of Plasmodium falciparum by Single-Molecule Long-Read Sequencing. Front Cell Infect Microbiol 11, 631545 (2021).

21. J. Dekker, K. Rippe, M. Dekker, N. Kleckner, Capturing chromosome conformation. Science 295, 1306–1311 (2002).

22. J. Dostie et al., Chromosome Conformation Capture Carbon Copy (5C): a massively parallel solution for mapping interactions between genomic elements. Genome Res 16, 1299–1309 (2006).

23. L. Cui, Q. Fan, L. Cui, J. Miao, Histone lysine methyltransferases and demethylases in Plasmodium falciparum. Int J Parasitol 38, 1083–1097 (2008).

24. J. R. Dixon et al., Topological domains in mammalian genomes identified by analysis of chromatin interactions. Nature 485, 376–380 (2012).

25. L. Dembele et al., Persistence and activation of malaria hypnozoites in long-term primary hepatocyte cultures. Nat Med 20, 307–312 (2014).

26. X. Deng et al., Bipartite structure of the inactive mouse X chromosome. Genome Biol 16, 152 (2015).

27. E. M. Bunnik et al., Comparative 3D genome organization in apicomplexan parasites. Proc Natl Acad Sci U S A 116, 3183–3192 (2019).

28. G. Batugedara et al., The chromatin bound proteome of the human malaria parasite. Microb Genom 6, (2020).

29. Z. J. Lorkovic, MORC proteins and epigenetic regulation. Plant Signal Behav 7, 1561–1565 (2012).

30. D. Q. Li, S. S. Nair, R. Kumar, The MORC family: new epigenetic regulators of transcription and DNA damage response. Epigenetics 8, 685–693 (2013).

31. N. E. Weiser et al., MORC-1 Integrates Nuclear RNAi and Transgenerational Chromatin Architecture to Promote Germline Immortality. Dev Cell 41, 408–423 e407 (2017).

32. S. Singh et al., The PfAP2-G2 transcription factor is a critical regulator of gametocyte maturation. Mol Microbiol 115, 1005–1024 (2021).

33. D. C. Farhat et al., A MORC-driven transcriptional switch controls Toxoplasma developmental trajectories and sexual commitment. Nat Microbiol 5, 570–583 (2020).

34. H. Wang, L. Zhang, Q. Luo, J. Liu, G. Wang, MORC protein family-related signature within human disease and cancer. Cell Death Dis 12, 1112 (2021).

35. Z. Zhong et al., MORC proteins regulate transcription factor binding by mediating chromatin compaction in active chromatin regions. Genome Biol 24, 96 (2023).

36. N. Saksouk et al., Histone-modifying complexes regulate gene expression pertinent to the differentiation of the protozoan parasite Toxoplasma gondii. Mol Cell Biol 25, 10301–10314 (2005).

37. A. Bougdour et al., Drug inhibition of HDAC3 and epigenetic control of differentiation in Apicomplexa parasites. J Exp Med 206, 953–966 (2009).

38. C. Hillier et al., Landscape of the Plasmodium Interactome Reveals Both Conserved and Species-Specific Functionality. Cell Rep 28, 1635–1647 e1635 (2019).

39. M. D. Jeninga, J. E. Quinn, M. Petter, ApiAP2 Transcription Factors in Apicomplexan Parasites. Pathogens 8, (2019).

40. T. Hiyoshi, J. A. Wada, Feline amygdaloid kindling and the sleep-waking pattern: observations on daily 22-hour polygraphic recording. Epilepsia 31, 131–138 (1990).

41. M. K. Singh et al., A nuclear protein, PfMORC confers melatonin dependent synchrony of the human malaria parasite P. falciparum in the asexual stage. Sci Rep 11, 2057 (2021).

42. A. K. Subudhi et al., DNA-binding protein PfAP2-P regulates parasite pathogenesis during malaria parasite blood stages. Nat Microbiol 8, 2154–2169 (2023).

43. J. M. Bryant et al., Exploring the virulence gene interactome with CRISPR/dCas9 in the human malaria parasite. Mol Syst Biol 16, e9569 (2020).

44. B. V. A. Singh Maneesh Kumar, Da Silva Israel Tojal, Santiago Verônica Feijoli, Moraes Miriam S., Adderley Jack, Doerig Christian, Palmisano Giuseppe, Llinás Manuel, Garcia Célia R. S., A Plasmodium falciparum MORC protein complex modulates epigenetic control of gene expression through interaction with heterochromatin eLIfe 12, (2023).

45. S. M. Ganesan, A. Falla, S. J. Goldfless, A. S. Nasamu, J. C. Niles, Synthetic RNA-protein modules integrated with native translation mechanisms to control gene expression in malaria parasites. Nat Commun 7, 10727 (2016).

46. M. Filarsky et al., GDV1 induces sexual commitment of malaria parasites by antagonizing HP1-dependent gene silencing. Science 359, 1259–1263 (2018).

47. M. Yuda, I. Kaneko, Y. Murata, S. Iwanaga, T. Nishi, Mechanisms of triggering malaria gametocytogenesis by AP2-G. Parasitol Int 84, 102403 (2021).

48. R. S. Kent et al., Inducible developmental reprogramming redefines commitment to sexual development in the malaria parasite Plasmodium berghei. Nat Microbiol 3, 1206–1213 (2018).

49. H. Choi, S. Kim, D. Fermin, C. C. Tsou, A. I. Nesvizhskii, QPROT: Statistical method for testing differential expression using protein-level intensity data in label-free quantitative proteomics. J Proteomics 129, 121–126 (2015).

50. W. A. M. Hoeijmakers et al., Epigenetic reader complexes of the human malaria parasite, Plasmodium falciparum. Nucleic Acids Res 47, 11574–11588 (2019).

51. C. Flueck et al., A major role for the Plasmodium falciparum ApiAP2 protein PfSIP2 in chromosome end biology. PLoS Pathog 6, e1000784 (2010).

52. X. Shang et al., Genome-wide landscape of ApiAP2 transcription factors reveals a heterochromatin-associated regulatory network during Plasmodium falciparum blood-stage development. Nucleic Acids Res 50, 3413–3431 (2022).

53. Q. Chen et al., Developmental selection of var gene expression in Plasmodium falciparum. Nature 394, 392–395 (1998).

54. A. Scherf et al., Antigenic variation in malaria: in situ switching, relaxed and mutually exclusive transcription of var genes during intra-erythrocytic development in Plasmodium falciparum. EMBO J 17, 5418–5426 (1998).

55. L. Meerstein-Kessel et al., Probabilistic data integration identifies reliable gametocyte-specific proteins and transcripts in malaria parasites. Sci Rep 8, 410 (2018).

56. M. Zhang et al., Uncovering the essential genes of the human malaria parasite Plasmodium falciparum by saturation mutagenesis. Science 360, (2018).

57. S. J. Goldfless, J. C. Wagner, J. C. Niles, Versatile control of Plasmodium falciparum gene expression with an inducible protein-RNA interaction. Nat Commun 5, 5329 (2014).

58. A. S. Nasamu et al., An integrated platform for genome engineering and gene expression perturbation in Plasmodium falciparum. Sci Rep 11, 342 (2021).

59. T. Kiss, Small nucleolar RNAs: an abundant group of noncoding RNAs with diverse cellular functions. Cell 109, 145–148 (2002).

60. N. Servant et al., HiC-Pro: an optimized and flexible pipeline for Hi-C data processing. Genome Biol 16, 259 (2015).

61. E. M. Bunnik et al., Changes in genome organization of parasite-specific gene families during the Plasmodium transmission stages. Nat Commun 9, 1910 (2018).

62. A. R. Ardakany, F. Ay, S. Lonardi, Selfish: discovery of differential chromatin interactions via a self-similarity measure. Bioinformatics 35, i145–i153 (2019).

63. N. Varoquaux, F. Ay, W. S. Noble, J. P. Vert, A statistical approach for inferring the 3D structure of the genome. Bioinformatics 30, i26–33 (2014).

64. A. H. Tencer et al., Molecular mechanism of the MORC4 ATPase activation. Nat Commun 11, 5466 (2020).

65. Y. Shao et al., Involvement of histone deacetylation in MORC2-mediated down-regulation of carbonic anhydrase IX. Nucleic Acids Res 38, 2813–2824 (2010).

66. F. He et al., Structural insight into the zinc finger CW domain as a histone modification reader. Structure 18, 1127–1139 (2010).

67. Y. Zhang et al., MORC3 Forms Nuclear Condensates through Phase Separation. iScience 17, 182–189 (2019).

68. J. Luo, S. Zeng, C. Tian, MORC4 Promotes Chemoresistance of Luminal A/B Breast Cancer via STAT3-Mediated MID2 Upregulation. Onco Targets Ther 13, 6795–6803 (2020).

69. Y. Mimura, K. Takahashi, K. Kawata, T. Akazawa, N. Inoue, Two-step colocalization of MORC3 with PML nuclear bodies. J Cell Sci 123, 2014–2024 (2010).

70. W. Trager, J. B. Jensen, Human malaria parasites in continuous culture. 1976. J Parasitol 91, 484-486 (2005).

71. D. A. Fidock, T. Nomura, T. E. Wellems, Cycloguanil and its parent compound proguanil demonstrate distinct activities against Plasmodium falciparum malaria parasites transformed with human dihydrofolate reductase. Mol Pharmacol 54, 1140–1147 (1998).

72. K. Rajaram, H. B. Liu, S. T. Prigge, Redesigned TetR-Aptamer System To Control Gene Expression in Plasmodium falciparum. mSphere 5, (2020).

73. L. Florens, M. P. Washburn, Proteomic analysis by multidimensional protein identification technology. Methods Mol Biol 328, 159–175 (2006).

74. T. Xu et al., ProLuCID: An improved SEQUEST-like algorithm with enhanced sensitivity and specificity. J Proteomics 129, 16–24 (2015).

75. D. L. Tabb, W. H. McDonald, J. R. Yates, 3rd, DTASelect and Contrast: tools for assembling and comparing protein identifications from shotgun proteomics. J Proteome Res 1, 21–26 (2002).

76. Y. Zhang, Z. Wen, M. P. Washburn, L. Florens, Refinements to label free proteome quantitation: how to deal with peptides shared by multiple proteins. Anal Chem 82, 2272–2281 (2010).

77. H. Choi, D. Fermin, A. I. Nesvizhskii, Significance analysis of spectral count data in label-free shotgun proteomics. Mol Cell Proteomics 7, 2373–2385 (2008).

78. Y. Zhang et al., Model-based analysis of ChIP-Seq (MACS). Genome Biol 9, R137 (2008).

79. M. K. Gupta, T. Lenz, K. G. Le Roch, Chromosomes Conformation Capture Coupled with Next-Generation Sequencing (Hi-C) in Plasmodium falciparum. Methods Mol Biol 2369, 15–25 (2021).

80. T. Yang et al., HiCRep: assessing the reproducibility of Hi-C data using a stratum-adjusted correlation coefficient. Genome Res 27, 1939–1949 (2017).

81. F. Ay, T. L. Bailey, W. S. Noble, Statistical confidence estimation for Hi-C data reveals regulatory chromatin contacts. Genome Res 24, 999–1011 (2014).

82. T. D. Goddard et al., UCSF ChimeraX: Meeting modern challenges in visualization and analysis. Protein Sci 27, 14–25 (2018).

