## Supplementary figures and images for "*Pf*MORC protein regulates chromatin accessibility and transcriptional repression in the human malaria parasite, *Plasmodium falciparum*"

### Figure S1

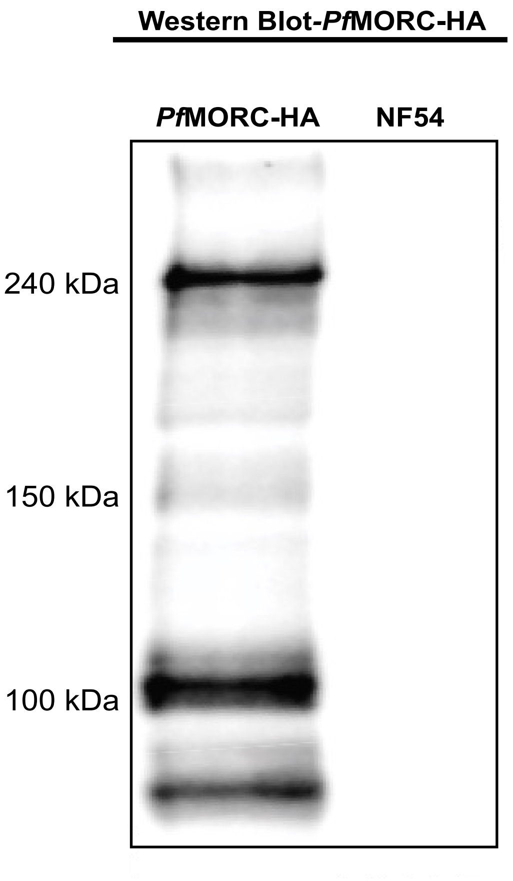

### Figure S2

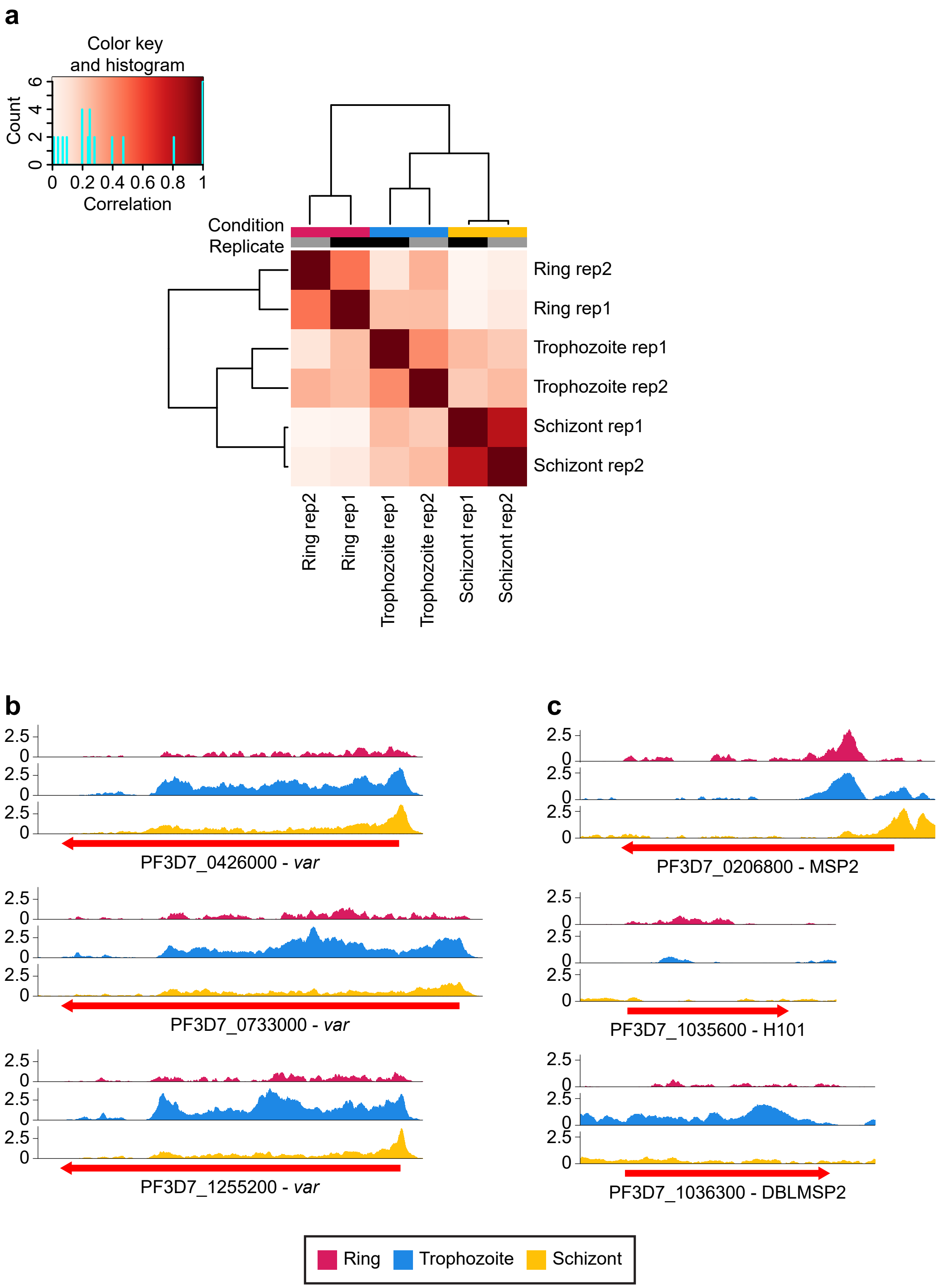

### Figure S3

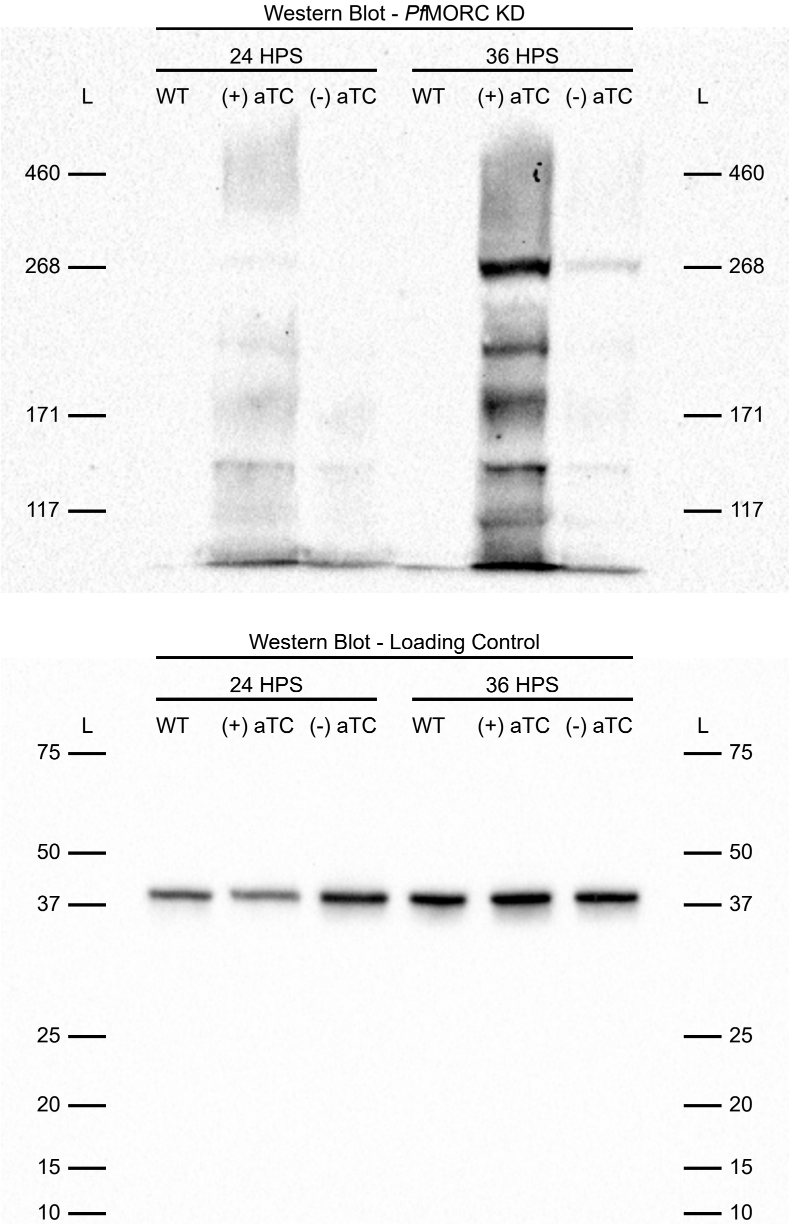

### Figure S4

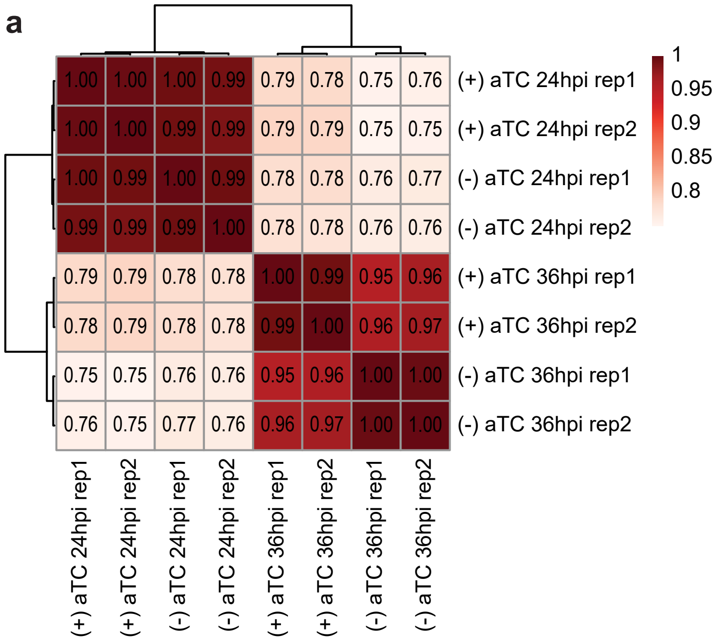

### Figure S5

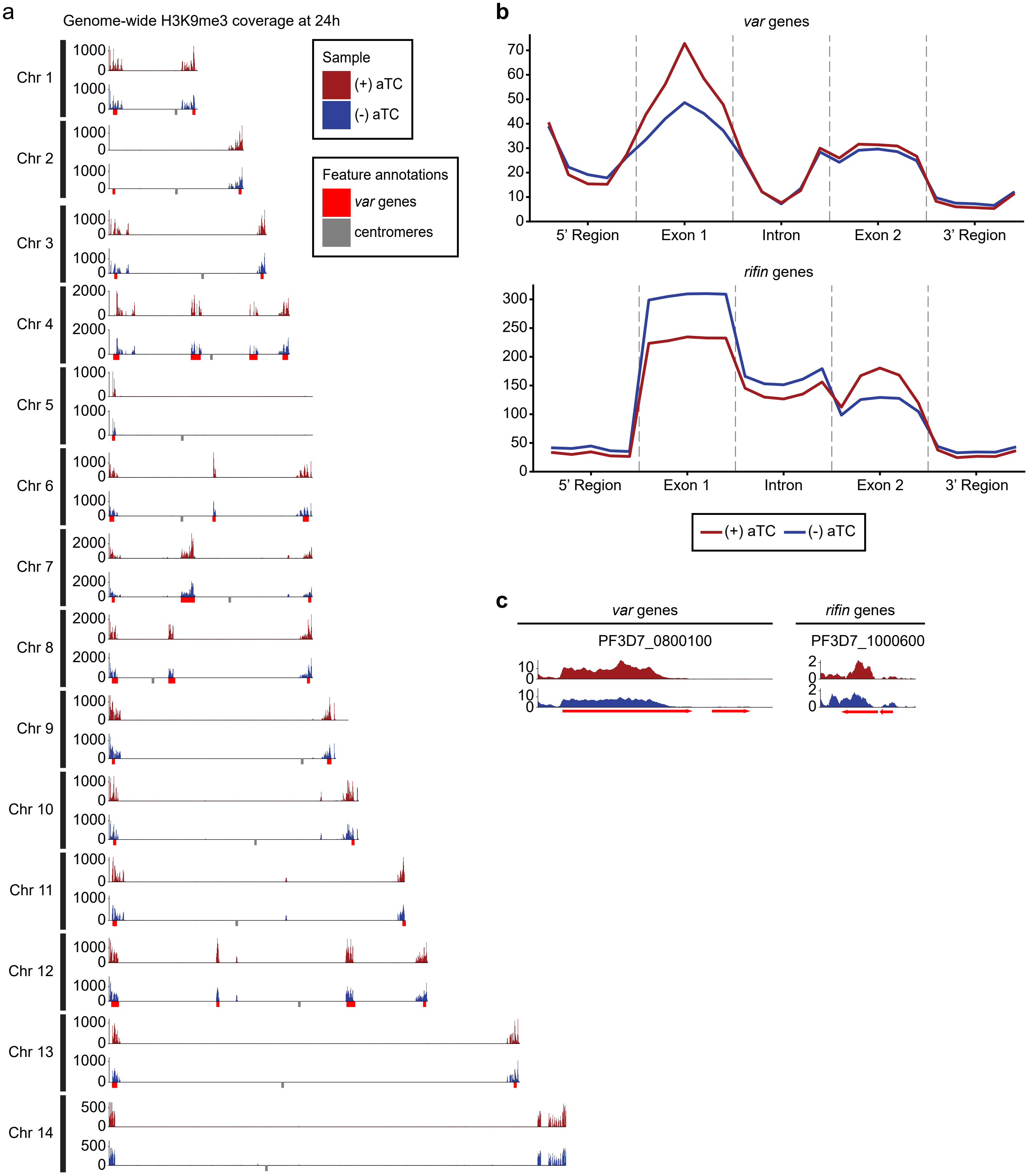

### Figure S6

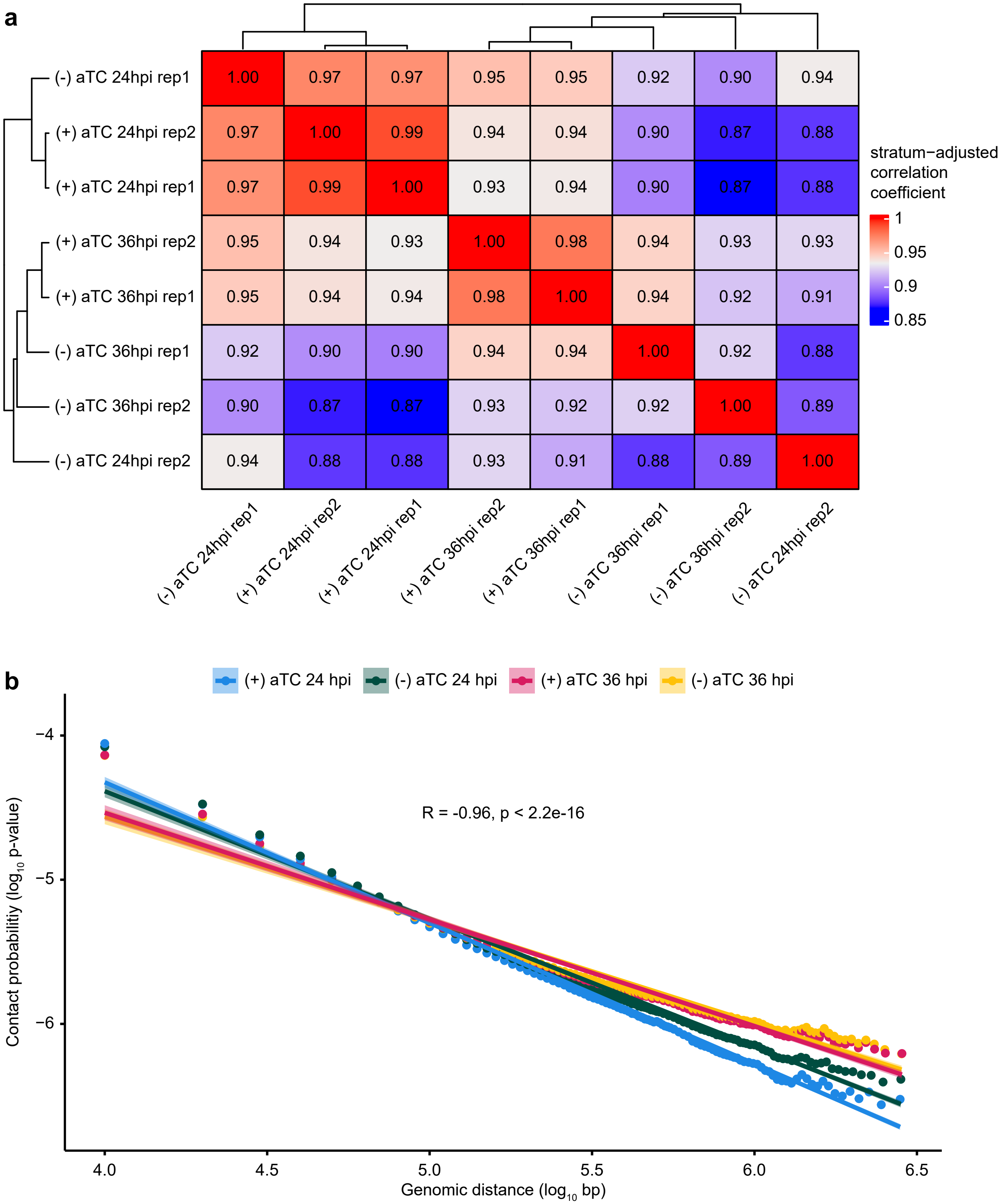

### Figure S7

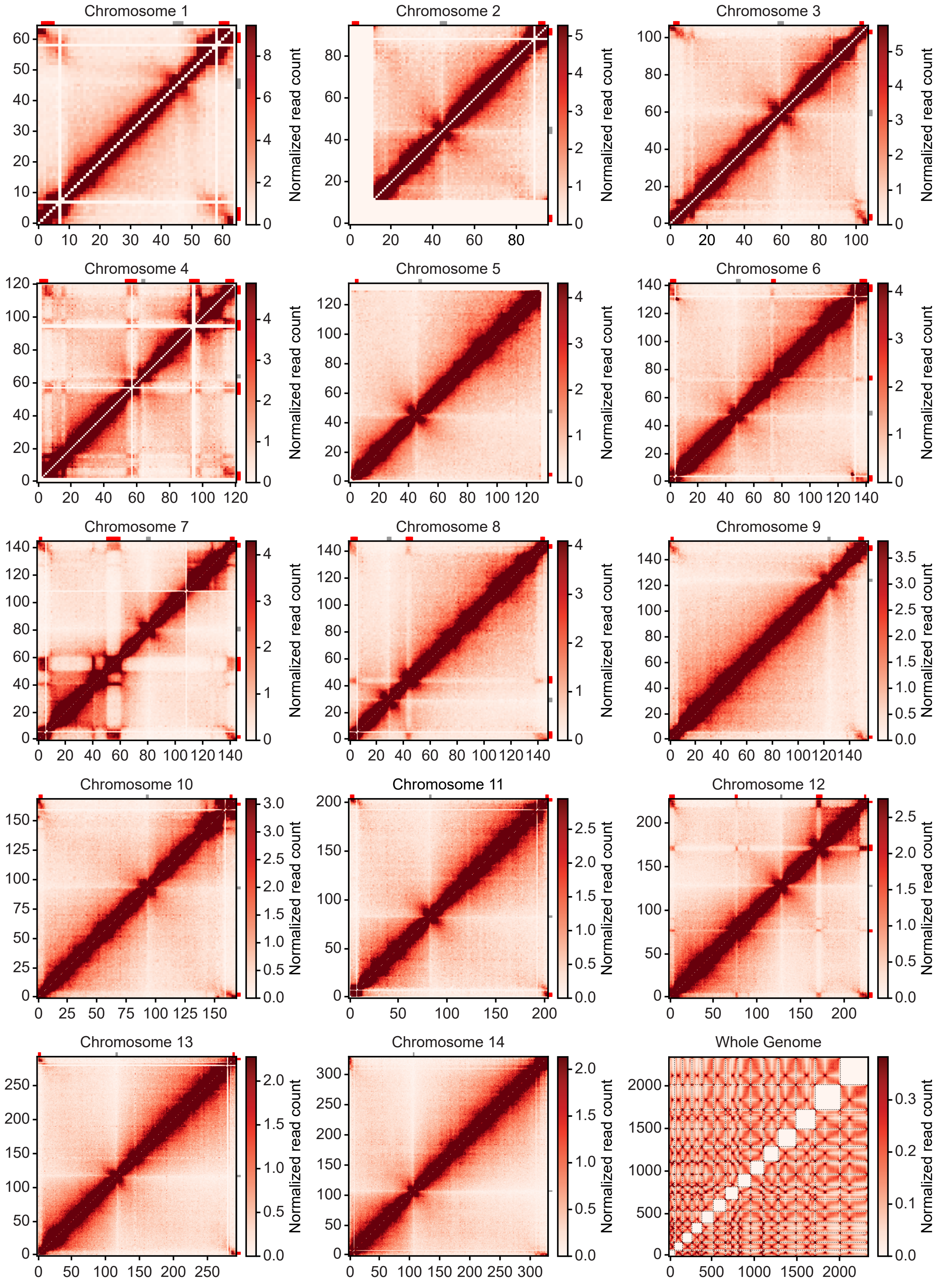

### Figure S8

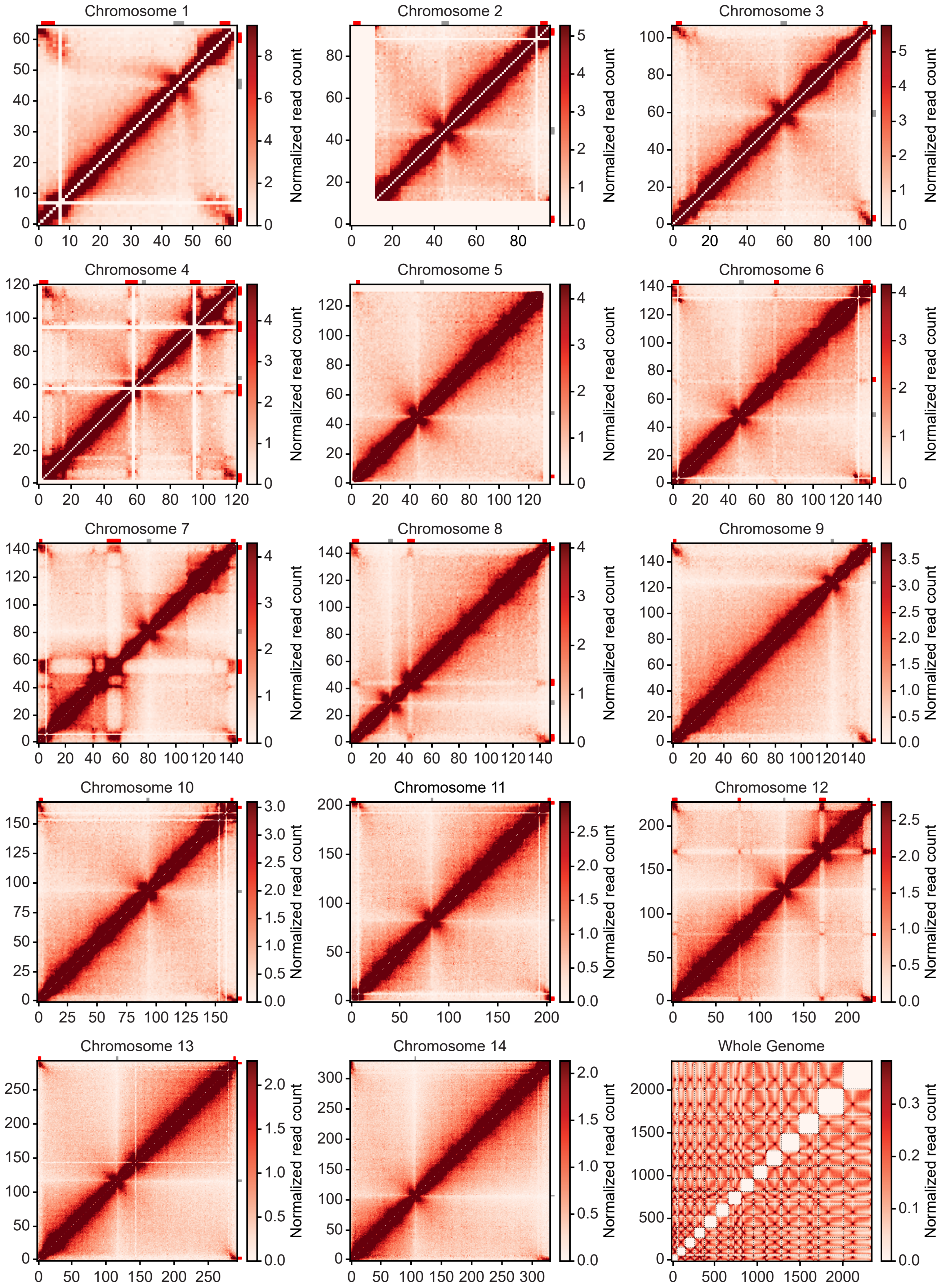

### Figure S9

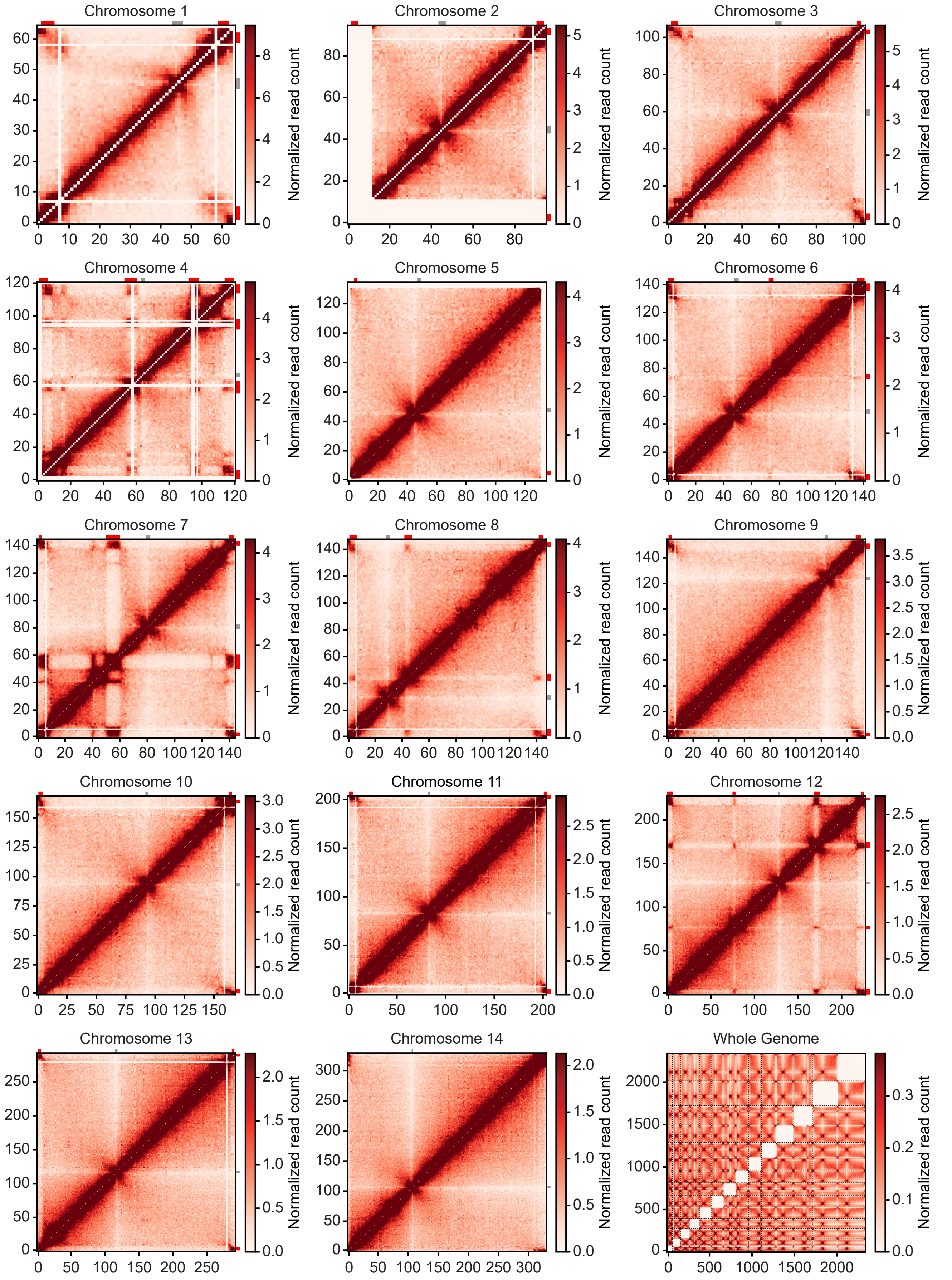

### Figure S10

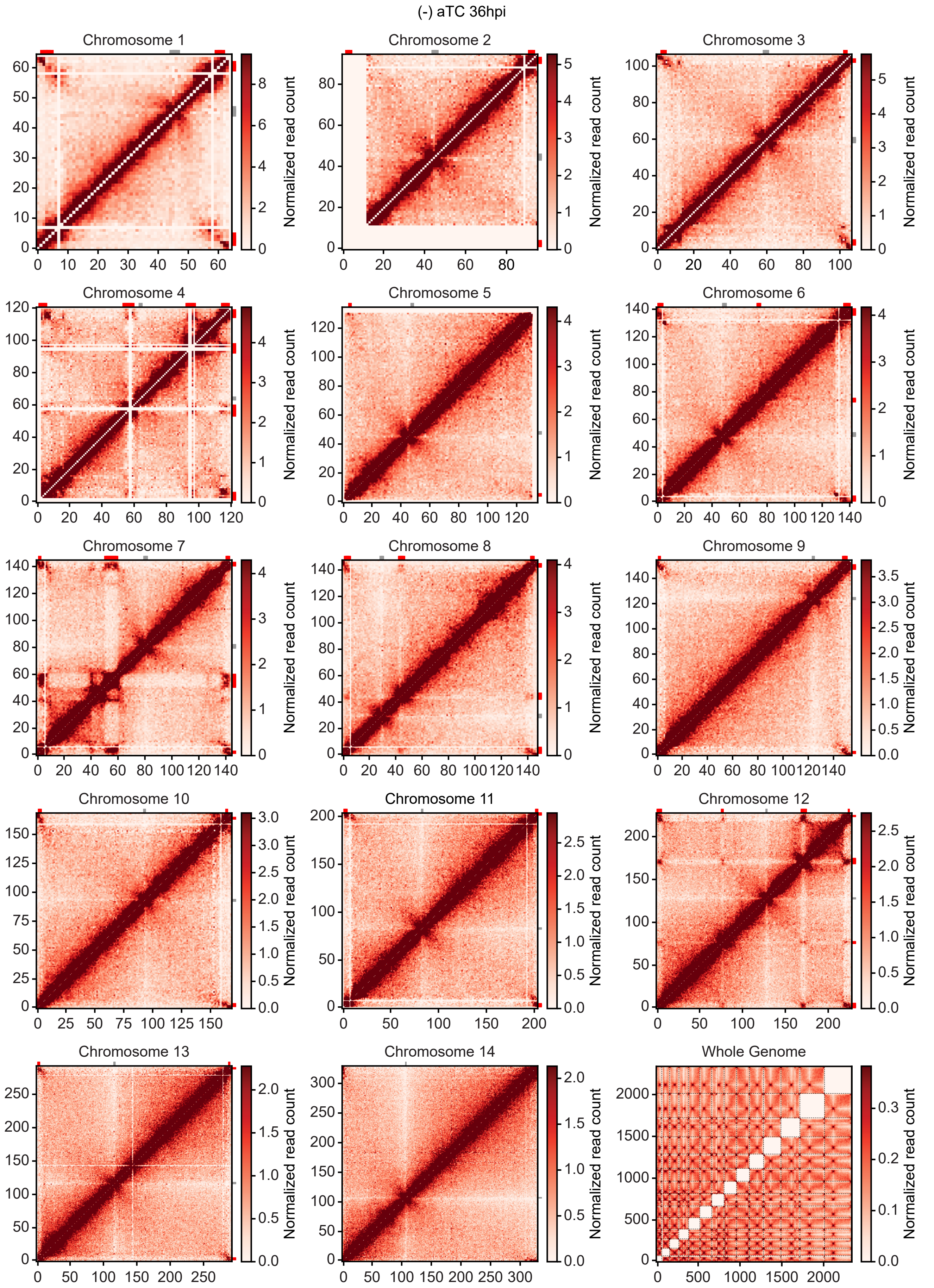

### Figure S11

(+) aTC 24hpi vs (-) aTC 24hpi differential interactions

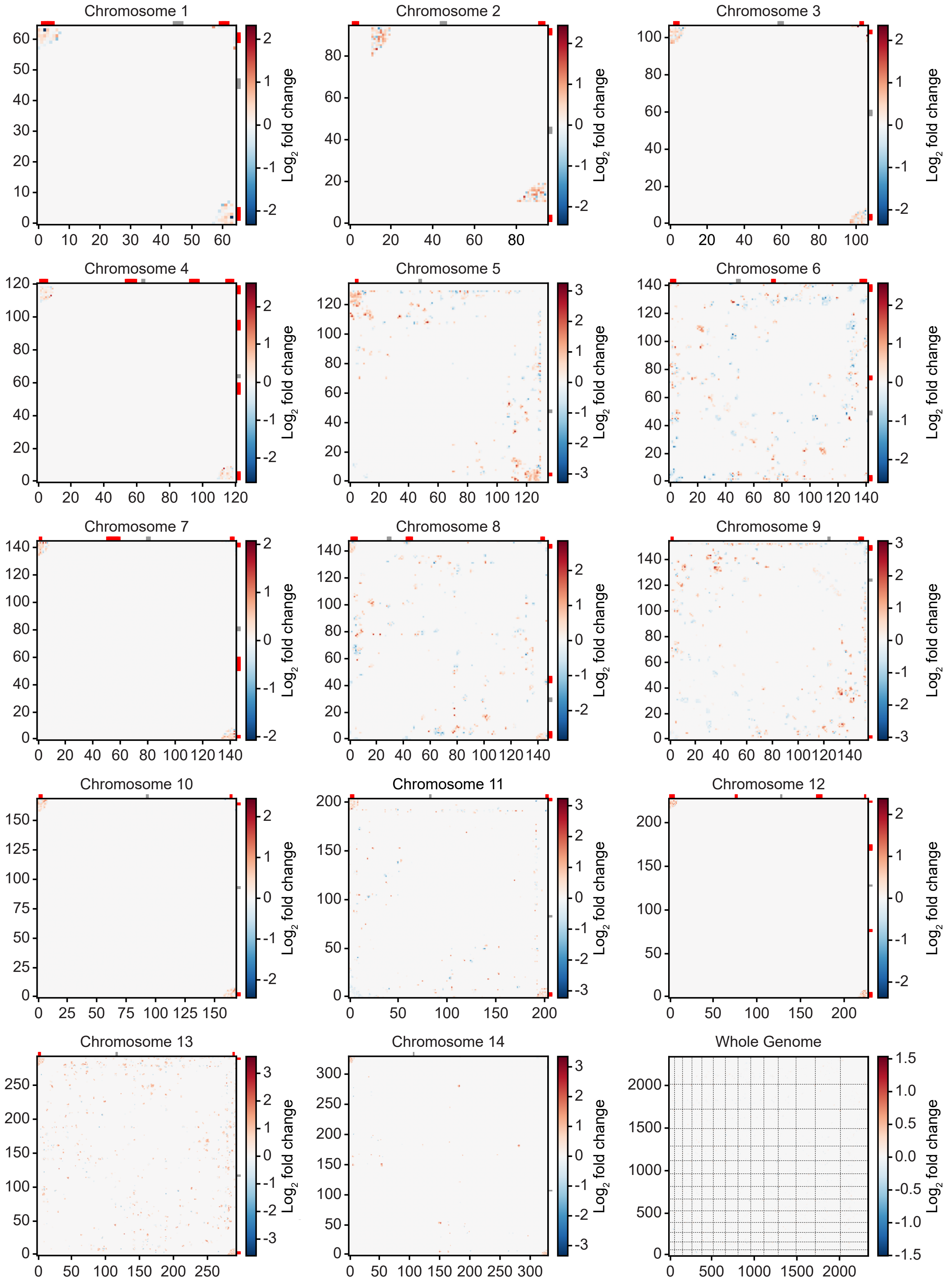

### Figure S12

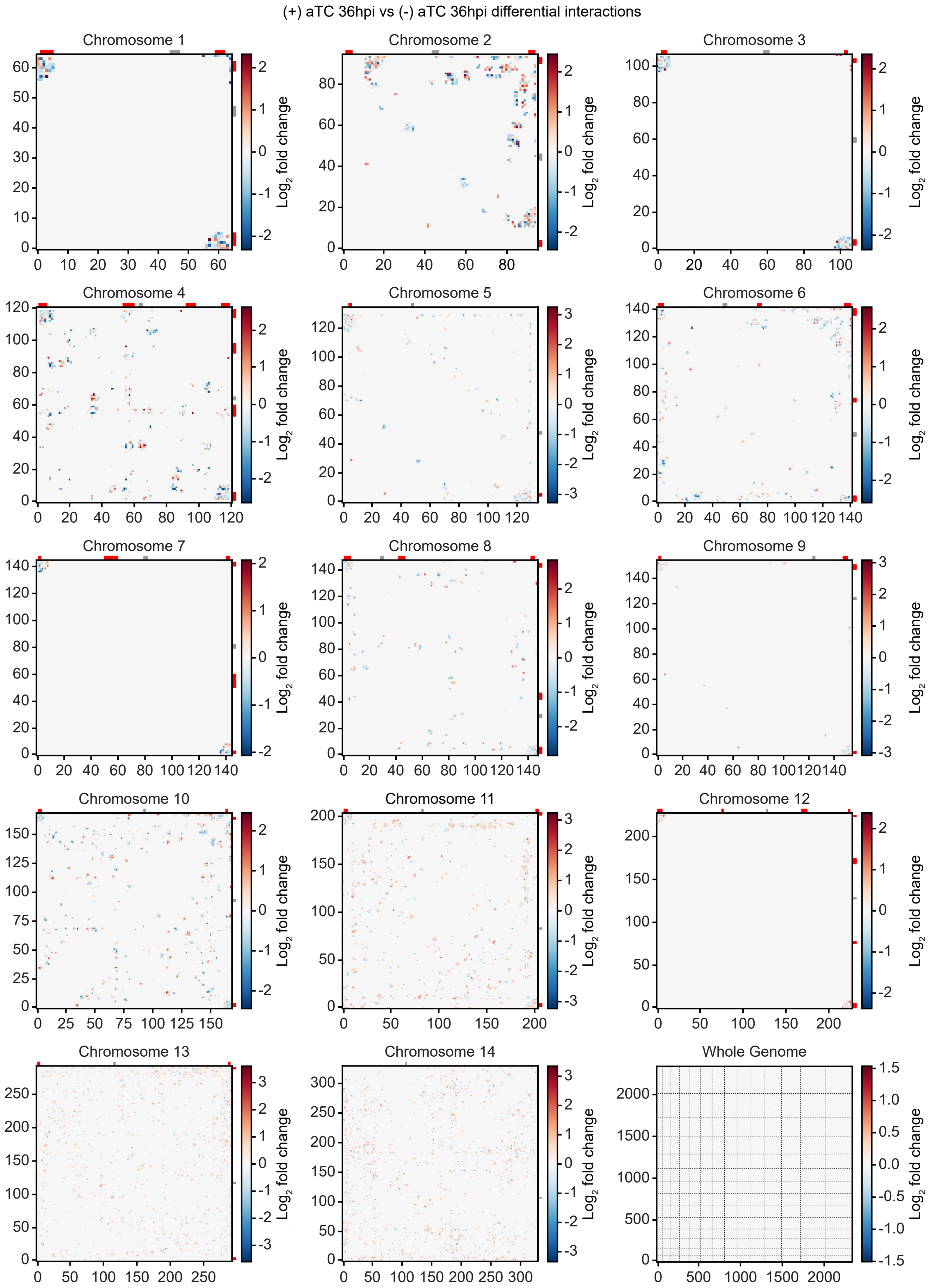
